# Predicting patient treatment response and resistance via single-cell transcriptomics of their tumors

**DOI:** 10.1101/2022.01.11.475728

**Authors:** Sanju Sinha, Rahulsimham Vegesna, Saugato Rahman Dhruba, Wei Wu, D. Lucas Kerr, Oleg V. Stroganov, Ivan Grishagin, Kenneth D. Aldape, Collin M. Blakely, Peng Jiang, Craig J. Thomas, Trever G. Bivona, Alejandro A. Schäffer, Eytan Ruppin

## Abstract

Tailoring the best treatments to cancer patients is an important open challenge. Here, we build a precision oncology data science and software framework for <u>PER</u>sonalized single-<u>C</u>ell <u>E</u>xpression-based <u>P</u>lanning for <u>T</u>reatments <u>In On</u>cology (PERCEPTION). Our approach capitalizes on recently published matched bulk and single-cell transcriptome profiles of large-scale cell-line drug screens to build treatment response models from patients’ single-cell (SC) tumor transcriptomics. First, we show that PERCEPTION successfully predicts the response to monotherapy and combination treatments in screens performed in cancer and patient-tumor-derived primary cells based on SC-expression profiles. Second, it successfully stratifies responders to combination therapy based on the patients’ tumor’s SC-expression in two very recent multiple myeloma and breast cancer clinical trials. Thirdly, it captures the development of clinical resistance to five standard tyrosine kinase inhibitors using tumor SC-expression profiles obtained during treatment in a lung cancer patients’ cohort. Notably, PERCEPTION outperforms state-of-the-art bulk expression-based predictors in all three clinical cohorts. In sum, this study provides a first-of-its-kind conceptual and computational method that is predictive of response to therapy in patients, based on the clonal SC gene expression of their tumors.

## Introduction

Precision oncology has made important strides in advancing cancer patient treatment in recent years, as reviewed in (Tsimberidou et al. 2020a; Huang et al. 2021; Bhinder et al. 2021; Singla and Singla 2020; Senft et al. 2017; Tsimberidou et al. 2020b). Much of the focus in the field has been on efforts to use recent FDA-approved NGS panels to identify “actionable” mutations in cancer driver genes, to match patients to treatments (Tsimberidou et al. 2020a). These efforts have been further boosted by the recent progress made in DNA-based liquid biopsies, which further can help guide and monitor treatment (Siravegna et al. 2017; Heitzer et al. 2019; Sawabata 2020). However, a large fraction of cancer patients still do not benefit from such targeted therapies, and efforts are hence much needed to find ways to analyze other molecular omics data types to benefit more patients. Addressing this challenge, recent studies have begun to explore the benefit of collecting and analyzing bulk tumor transcriptomics data to guide cancer patient treatment (Beaubier et al., 2019; Hayashi et al., 2020; Rodon et al., 2019; Tanioka et al., 2018; Vaske et al., 2019; Wong et al., 2020, Lee et al., 2021). These studies have demonstrated the potential of such approaches to complement DNA sequencing approaches in increasing the benefit from omics-guided precision treatments to patients.

One key limitation of current genomic and transcriptomic treatment approaches is that they are mostly based on bulk tumor data. Tumors are typically heterogeneous and composed of numerous clones, making treatments targeting multiple clones more likely to diminish the likelihood of resistance emerging due to clonal selection, and hence potentially enhancing the overall patient’s response (Castro et al. 2021). This fundamental challenge has been driving two major developments in recent years, the search for effective treatment combinations and the advent of single-cell profiling of the tumor and its microenvironment.

Briefly, large-scale combinatorial pharmacological screens have been performed in patient-derived primary cells, xenografts, and organoids and have already given rise to numerous combination treatment candidates (e.g., Wensink, et al. 2021, Yao et al. 2020, de Witte et al. 2020). Concomitantly, the characterization of the tumor microenvironment via single-cell omics has already led to important insights regarding the complex network of tumor-microenvironment interactions involving both stromal and immune cell types (Castro et al. 2021). It also offers a promising way to learn and predict drug response at a single-cell resolution. The latter, if successful, could guide the design of drug treatments that target multiple tumor clones disjointly (Shalek et al 2017, Adam et al 2020, Zhu et al 2017) and help us understand the ensuing resistance to better overcome it. However, building such predictors of drug response at a single cell (SC) resolution is currently challenging due to the paucity of large-scale preclinical or clinical training datasets. Previous efforts, including a recent computational method - *Beyondcell*, that identifies tumor cell subpopulations with distinct drug responses from single-cell RNA-seq data for proposing cancer-specific treatments, have focused on preclinical models and lacks validation in patients at the clinical level (Kim et al 2016, Suphavilai et al 2020, Fustero-Torre et al. 2021). Additional efforts to identify biomarkers of response and resistance at the patient level using single-cell expression are rapidly emerging for both targeted- and immuno-therapies, with remarkable results (Cohen et al 2021, Ledergor et al 2018, Sade-Feldman et al 2018). However, to date, the feasibility of harnessing SC patients’ tumor transcriptomics for tailoring patients’ treatment in a direct, systematic manner has yet to be described.

Aiming to address this challenge systematically, here we present a precision oncology framework for <u>PER</u>sonalized single-<u>C</u>ell <u>E</u>xpression-based <u>P</u>lanning for <u>T</u>reatments <u>In ON</u>cology (PERCEPTION). This approach builds upon the recent availability of large-scale pharmacological screens and SC expression data in cancer cell lines to build machine learning-based predictors of drug response based on the gene expression of single cells. We first show that the predicted viability for drugs with known mechanisms of action strongly correlates with the pathway activity it is targeting at single-cell resolution, visualizing our ability to predict at this resolution. Second, we show that the PERCEPTION can successfully predict the response to single and combination treatments in three independent screens performed in cancer and patient-tumor-derived primary cells based on their SC-expression profiles. Thirdly, we show that PERCEPTION successfully stratifies responders vs. non-responders to combination therapies in two recently published multiple myeloma and breast cancer clinical trials using their tumor SC-expression profile. Finally, we show that it also successfully captures the development of resistance and cross-resistance to four different kinase inhibitors in a recently published cohort of lung cancer patients with tumor SC-expression profiles during treatment. Notably, PERCEPTION outperforms state-of-the-art bulk-based predictors in all three SC clinical cohorts considered.

## Results

### Overview of PERCEPTION

To predict patient response to a therapy using their tumor’s SC-expression profile, we built a machine learning pipeline called PERCEPTION (**Figure 1A**, a detailed description is provided in **Methods**). PERCEPTION builds drug response models from large-scale pharmacological screens performed in cancer cell lines where *both* bulk and SC-expression are available. As there is currently a paucity of large-scale matched response and single-cell data either in pre-clinical studies or in patients, we designed a stepwise prediction pipeline: first, the prediction models are trained on large-scale bulk-expression profiles of cancer cell lines and then, in a second step, the models’ performance is further optimized by training on SC-expression profiles of cancer cell lines. To this end, we mined bulk-expression (Ghandi et al 2020) and drug response profiles (PRISM) of 488 cancer cell lines (**Table S1**) from the DepMap database (Tsherniak. et al 2017). The SC-expression profiles of these cell lines (N=205, **Table S1**) have been obtained from Kinker et al. 2020. Drug efficacy is measured via area under the curve (AUC) viability-dosage curve, where lower AUC values indicate increased sensitivity to treatment (**Table S1**). We note that, due to a lack of large-scale screens in normal and immune cells for training, we could only train our drug response models on cancer cells in this study.

**Figure 1.**
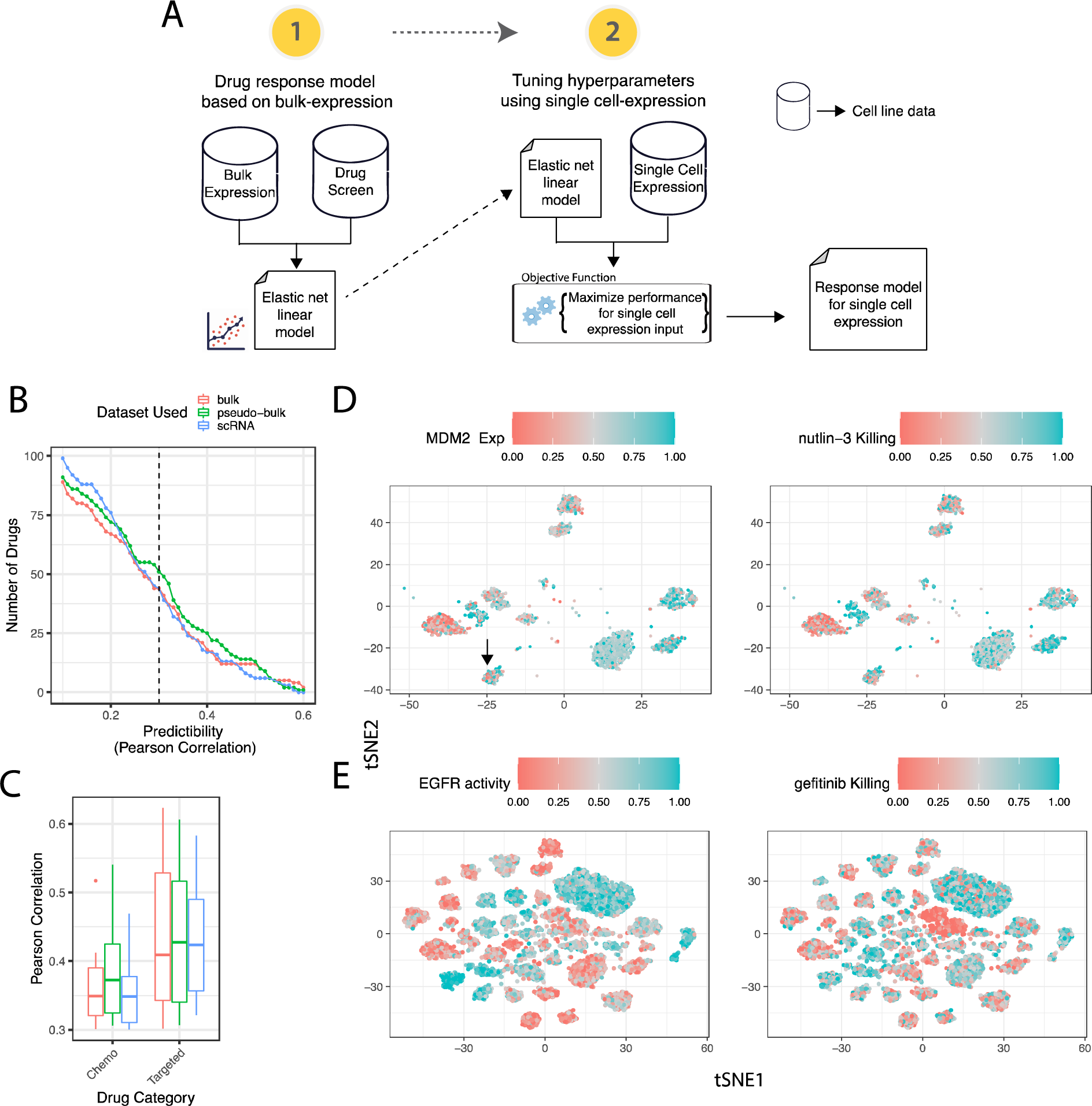
Overview of PERCEPTION framework. (**A**) <u>Building PERCEPTION prediction models is performed in two steps:</u> (i) Build response models based on drug response data measured in large-scale drug screens performed on cancer cell lines and their matched bulk expression. (ii) Tune these models by determining the optimal number of genes used as predictive features that maximize its prediction performance based on SC-expression of cancer cell lines. The mean predicted response over all those individual cells from a given cell line is taken as the predicted SC-based response of that cell line (Methods). **(B)** The number of predictive models (defined by Pearson R>0.3) for FDA-approved drugs for cancer generated by PERCEPTION, as tested via cross-validation (y-axis) when SC-expression (blue), bulk-expression (red), and pseudo-bulk are used for a Pearson correlation threshold (x-axis, Predictability). **(C)** The distribution of predictive performance (x-axis) of the predictive models. In the boxplots, the center line, box edges, and whiskers denote the median, interquartile range, and the rest of the distribution, respectively, except for points that were determined to be outliers using a method that is a function of the interquartile range, as in standard box plots. (**D)** In the left-most panel, visualizing the predicted killing by Nutlin-3, a canonical MDM2 antagonist, the predicted killing and the expression of MDM2 are provided for every single cell (each point) in the top and bottom tSNE plot, respectively. The intensity of the color denotes the extent of predicted killing in the left panel and measured MDM2 expression in the right pane, where the respective legends are provided. In this panel, we provide 3566 single-cells from nine p53 WT lung cancer cell lines. The tSNE clustering is performed using the expression profile of all the genes. **(E)** A similar display visualizes the predicted killing and the activity level of an EGFR pathway signature across 12482 individual lung cancer cells.

To build a predictor of response for a given drug, PERCEPTION performs the following two steps: **1**. It first builds a regularized linear model of drug response using the bulk expression and drug response data available for ∼300 cancer cell lines. **2**. In the second step, we determine the number of genes used as predictive features (hyperparameter tuning) that maximize its ability to predict the response from the SC-expression data, analyzing the ∼160 cancer cell lines for which there is also additional SC-expression data. The goal of this step is to enable PERCEPTION models to take SC-expression as input to predict drug response. To evaluate the performance of an SC model in a given cell line, PERCEPTION predicts the response to a given drug for each of its individual cells, and the mean response over all those individual cells is taken as the predicted SC-based response of that cell line to that specific drug. The output of this machine learning pipeline is hence a drug-specific response model and a quantification of its predictive accuracy from SC-expression, which importantly is evaluated in unseen test fraction of the cells, employing a standard leave-one-out (one cell line) cross-validation procedure (**Methods**).

### Building PERCEPTION models for FDA approved cancer drugs based on the PRISM screen

We applied PERCEPTION aiming to build response models for 133 U.S FDA-approved oncology drugs available in the drug screen PRISM (**Table S2**). The predictive performances for these drugs are provided in **Figure 1B**. We denoted models as predictive at a single-cell resolution, if the Pearson correlation between their predicted (mean SC-response per cell line) vs. observed viability on the leave-one-out test data was greater than 0.3. This threshold was chosen as it corresponds to the mean cross-screen replicate correlation observed among three major pharmacological screens (average cross-platform correlation across GDSC, CTD & PRISM ∼ 0.30), as reported previously in the literature (Corsello et al. 2020). We were able to build such predictive models for 33% (44 out of 133 drugs, **Table S2**) of the drugs (**Figure 1B)**. Studying the predictive accuracy of these 44 predictive models in a cross-validation manner for different transcriptomics inputs, including SC, bulk, and pseudo-bulk-expression (generated by summing up the gene-mapped reads across single cells, Methods), we reassuringly find that the predictive performance of PERCEPTION for SC-expression as inputs is comparable to that performance obtained using bulk-expression or pseudo-bulk as inputs (**Figure 1C**).

### Visualization of PERCEPTION’s ability to predict viability at single-cell resolution in two case studies

To visualize PERCEPTION*’s* ability to predict cell killing at single-cell resolution, we examined its predicted killing for two drugs, where the pathway involved in the mechanism of action of the drug is well characterized (nutlin-3 and erlotinib). We applied the PERCEPTION pipeline described above to build SC-based predictors for these two drugs and studied them further, as follows.

The first case involves the canonical antagonist, nutlin-3, whose mechanism of killing involves the inhibition of the interaction between *MDM2* and the tumor suppressor p53; thus, *MDM2* high activity is a known response biomarker to nutlin-3 treatment (Arya et al. 2010). Via PERCEPTION, we built a response model for nutlin-3, where the correlation between the predicted and observed response on the test set was: R = 0.598, P=1.2E-16, (with *MDM2* expression being one of the top-ranked predictive features). Using this model, we predicted the post-nutlin-3 treatment killing for each of 3566 single cells across nine p53 wild-type lung cancer cell lines. Across these single-cells, the predicted killing and *MDM2* expression are strongly correlated across the individual cells screened (Pearson R= 0.50, P<2E-16, as visualized in **Figure 1D**), as expected. The figure also shows a few sub-clones that have predicted pre-existing nutlin-3 resistance (**Figure 1D-arrow highlight**).

In the second case, we performed a similar analysis to study and visualize PERCEPTION*’s* ability to predict the response to erlotinib (with model’s test performance of Pearson’s R= 0.50, P<1E-05). Erlotinib targets oncogenic, activating mutations of epidermal growth factor receptor (*EGFR*) and has been used to treat non-small cell lung cancer. We reassuringly find that the predicted killing of erlotinib and *EGFR* pathway activity (estimated via the mean expression of a published *EGFR* signature (Cheng et al. 2020)) are significantly correlated across individual cells (Pearson R= 0.34, P<2E-16), as visualized in **Figure 1E**. Similar findings with other *EGFR* inhibitors developed more recently (afatinib, icotinib, lapatinib, osimertinib) with even stronger correlations strength and other FDA-approved drugs with well-characterized mechanisms of action are provided in **Extended Figure 1**.

### Evaluation of PERCEPTION models built on PRISM in the GDSC and a large lung cancer drug screen

We next aimed to evaluate the performance of PERCEPTION models that are built using the PRISM screen and other large-scale cell-line screens, for which we have matching SC cell-lines data (Garnett et al. 2012, Nair et al. 2021). To this end, we first identified the drugs that are shared between the PRISM and GDSC screens (N=191, **Table S3**, quality control and model building steps in **Methods**). We were able to build PRISM-based PERCEPTION predictive models for 16 drugs that have been screened in both PRISM and GDSC, and that have a substantial positive correlation between their AUC values (Pearson R > 0.3 and p-value < 0.05 in all cell lines). In building these models we have left out 80 randomly selected test cell lines (with SC-expression) that were used to test the performance of the resulting PERCEPTION models in each of the two screens. As a starting point for this comparison, we note that the mean correlation between the *experimental* viabilities reported in GDSC vs. PRISM (screen concordance) across the 80 shared test cell lines was 0.44 (**Figure 2A**, green, Methods). For the same testing set, the mean correlations between the predicted vs. observed viabilities, using the SC-based models generated by PERCEPTION, are 0.38 in the PRISM dataset (on which they were built, **Figure 2A**, blue) but is still considerable in the GDSC screen, attaining a correlation of 0.28 there (**Figure 2A**, orange). As expected, the prediction performance of the PRISM-based models in the GDSC test set is correlated with the concordance between the experimentally measured drug’s viability profiles in the two screens (Pearson R=0.49, P=5.89E-02; **Figure 2B, Table S4**). Of note, as the range of predicted values is typically smaller than those observed in the screens (**Extended Figure 2**), we use scaled predicted AUC scores (z-score) in the further analyses reported below.

**Figure 2:**
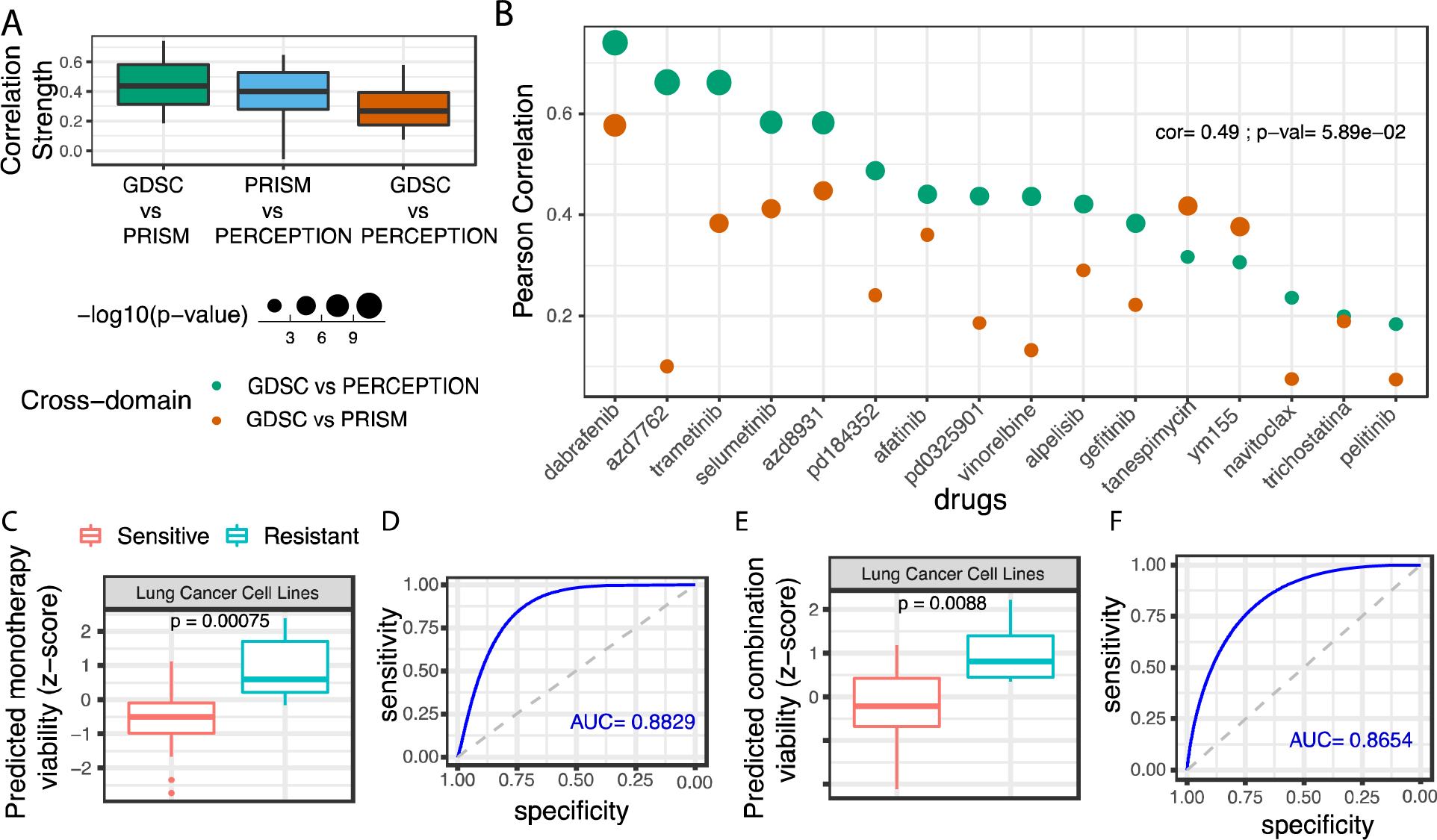
PERCEPTION’s performance in multiple cancer cell lines screens. **(A)** The correlations (Pearson Rho (y-axis)) comparing “GDSC vs. PRISM”, “PRISM vs. PERCEPTION” (cross-validation performance), and “GDSC vs. PERCEPTION”. Drug response predictions were performed at a single-cell resolution and the cell line level response (mean response across single cells) was used as the output prediction. **(B)** Relationship between the correlation between “GDSC vs. PRISM” experimental results (green) and the prediction accuracy of PRISM-based PERCEPTION models in prediction of the response in the GDSC screen (orange) across the 16 predictable drugs. The size of the dots represents the Pearson correlation-based p-value in -log10 scale. The drugs are ordered on the x-axis from left to right in the decreasing order of their correlation between GDSC and PERCEPTION responses. **C)** PERCEPTION predicted viability of monotherapies based on the cell lines SC-expression (x-axis) for resistant (N=72) vs. sensitive (N=84) cell lines, plotted via a standard boxplot. Significance is computed using a one-tailed Wilcoxon rank-sum test. **D)** A receiving operator curve is plotted showing the relationship between sensitivity and specificity, where the area under the curve denotes the power of stratification of sensitive vs resistant cell lines. The area under this curve is provided at the right corner. The area under the dashed diagonal line denotes a random-model performance. Panels **E)** and **F)**, respectively shows PERCEPTION’s ability to predict the response to drug combinations in that screens (28 resistant vs 24 sensitive cell lines).

Finally, we tested and validated the predictive power of PERCEPTION in another independent, yet unpublished drug screen in lung cancer cell lines (Nair et al. 2021), which includes monotherapies and combinations of 14 cancer drugs across 21 lung cancer cell lines in five dosages. As above, the PERCEPTION predictions are based on the SC data of the pertaining cell lines (shown in **Figure 2C-F**). For brevity, and since the main emphasis of our study is on building response predictors in patients, these validations and their results are described in more detail in the Supplementary material (**Extended Figure 3A-F, Table S5, Methods, Supplementary Note 1**). In sum, these analyses demonstrate PERCEPTION’s ability to predict drug monotherapy and combination response in independent cancer cell lines screens based on their SC-expression.

### SC-based PERCEPTION predictions in patient-derived Head and Neck cancer primary cells

Moving from cancer cell-line cells to patient tumor cells, we next tested the ability of PERCEPTION to predict response in patient-derived primary cells (PDC). We used SC-expression of head and neck cancer primary cells derived from five different patients treated with eight different drugs at two concentrations (**Table S6**), including both monotherapy and combination therapies (Suphavilai et al. 2020). We were able to build predictive PERCEPTION response models for 4 out of the 8 drugs tested (docetaxel, epothilone-b, gefitinib, and vorinostat; Pearson R threshold > 0.25, **Methods**) and focused our analysis on these drugs. For monotherapy treatments, the predicted viability is significantly higher in resistant vs. sensitive cell lines (top 40% vs. bottom 40% cell lines ranked by viability, N=8 each, **Figure 3A**), with an ROC-AUC of 0.64 (**Figure 3B**). The predicted viability over the 20 monotherapy-cell-lines combinations (4 monotherapies x 5 cell-lines) is correlated with the observed viability (Pearson R=0.46; P<0.03, **Extended Figure 4**), and individual drug-level correlations are provided in **Extended Figure 5**. Higher predicted viability in resistant cell lines is also observed for combination treatments (**Figure 3C**), with a ROC-AUC of 0.86 (**Figure 3D**). The predicted viability of combination treatment across 15 combination therapy-cell-line pairs (3 combinations x 5 cell lines) is highly correlated with the observed viability (Pearson R=0.73; P<0.002, **Extended Figure 4**). The predicted vs. experimental correlations obtained for all data points and drug levels are provided in **Extended Figures 4-5**. These results demonstrate the ability of PERCEPTION models to predict response in patient-derived single cells.

**Figure 3:**
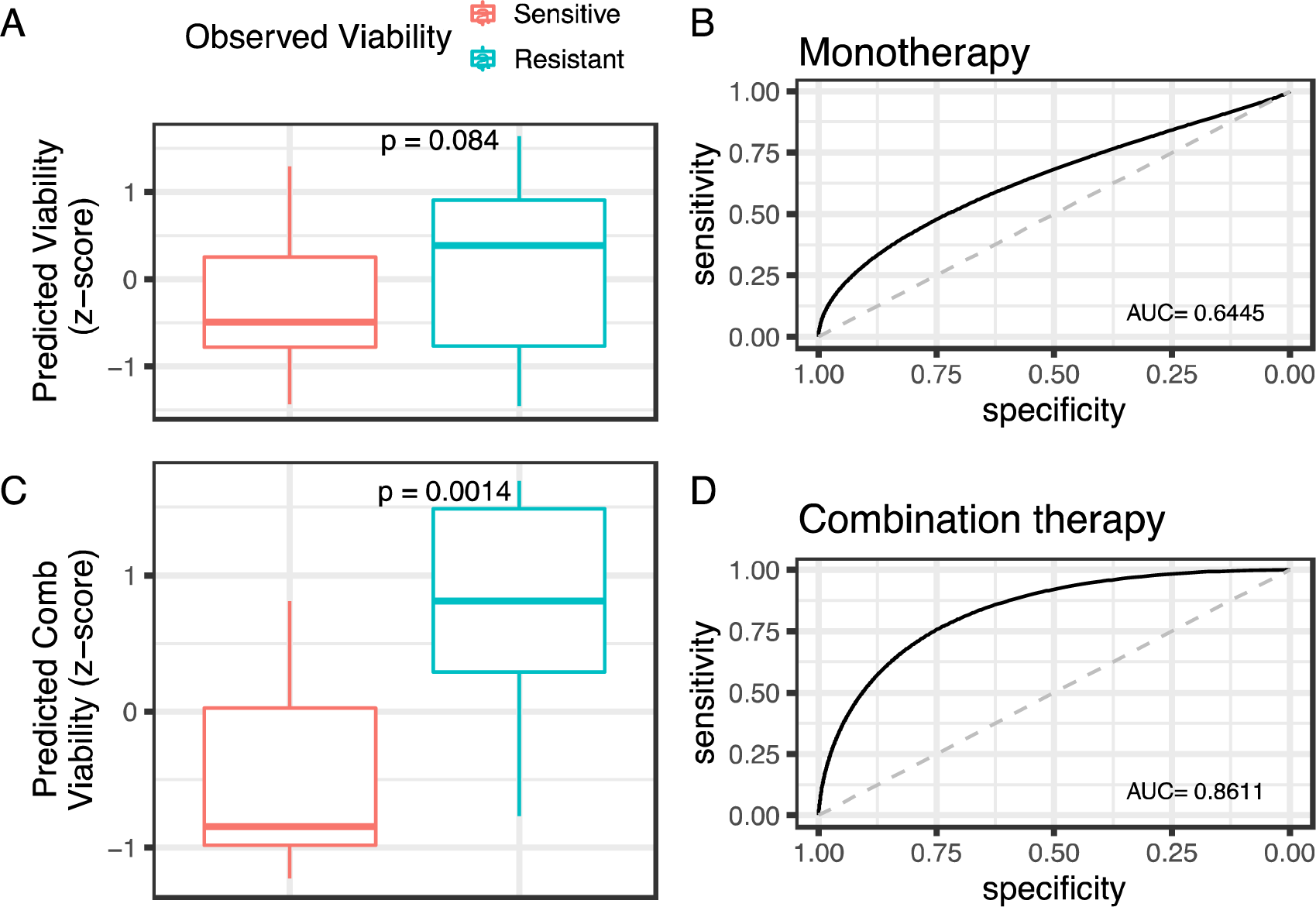
Prediction monotherapy and combination response in patient-derived primary cells. **(A)** PERCEPTION predicted viability in resistant (n=8) vs. sensitive (n=8) cell lines. **(B)** ROC curves depicting the prediction power (sensitivity and specificity) of the predicted viability to stratify resistant vs. sensitive cell lines. The area under this curve is provided at the right corner and denotes overall model prediction power. The area under the dashed diagonal line denotes a random-model performance. In **(C)** and **(D)**, we repeated the analysis for combination treatment (Number of resistant vs. sensitive cell lines=6 vs. 6). The boxplots provided are standard and the significances are computed using the one-tailed Wilcoxon rank-sum test.

### PERCEPTION predicts DARA-KRD based combination treatment response in a multiple myeloma clinical trial

We next turned to test the ability of *PERCEPTION* models to predict patient responses based on pre-treatment SC transcriptomics from their tumors, which is our main goal. Very few such datasets exist with considerable coverage of sequenced cancer cells and treatments involving drugs that PERCEPTION can currently predict. We began with the largest such dataset published to date, including data for 41 multiple myeloma patients. The patients were treated with a DARA–KRD combination of four drugs - daratumumab (monoclonal antibody targeting CD38), carfilzomib (proteasome inhibitor), lenalidomide (immunomodulator), and dexamethasone (anti-inflammatory corticosteroid) (Cohen et al. 2021). The cells were clustered based on their scRNA-seq profiles by (Cohen et al. 2021). The SC-expression and clonal (cluster) composition and treatment response labels were available for 28 tumor samples of these patients, whose pretreatment clonal composition as originally determined by (Cohen et al. 2021) is shown in **Figure 4C**. Patient response was measured via tumor size estimates in radiological images.

**Figure 4:**
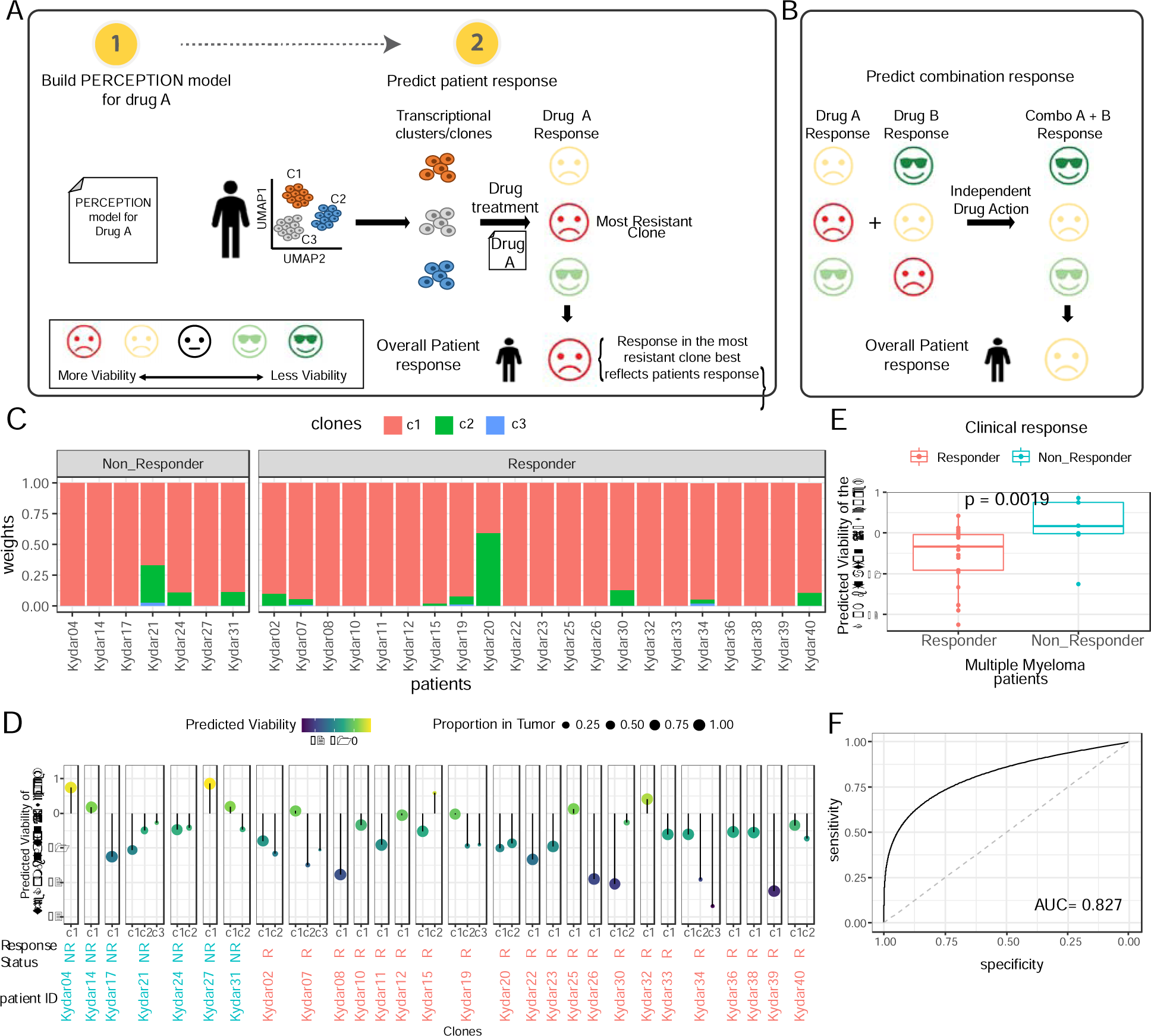
PERCEPTION stratifies responders vs. non-responders of the combination DACA-KRD therapy regime in multiple myeloma patients. **(A):** <u>Using PERCEPTION models to predict patient response</u> based on the clonal (transcriptional clusters) architecture of every tumor: **(i)** First, we generate PERCEPTION models for the drug A. **(ii)** Second, the scRNAseq of tumor cells in a patient is analyzed to identify the transcriptional clusters (or transcription-based clones) composition of each tumor. **(iii)** Third, we predict the response of the drug A separately for each cluster (the smiley faces represent the spectrum of drug response; sad - more viability or less killing to happy - less viability or more killing) – this prediction is done based on its mean expression. **(iv)** Finally, we consider the most resistant clone (clone with the highest viability) as the overall patient response. **(B)** In case of combination treatments, in the third step of **panel A**, we predict the response of each drug in a given combination separately for each cluster (the smiley faces represent the spectrum of drug response; sad - more viability or less killing to happy - less viability or more killing) – this prediction is done by taking the maximal killing effect observed for any of the drugs on that clone, based on its mean expression, as the overall response for that clone. This is motivated by the Independent Drug Action (IDA) principle, where it was shown that the predicted response of a combination of drugs is well represented by the effect of the single most effective drug in the combination (Ling et al. 2020). **(C)** Distribution of abundance of malignant sub-clones (y-axis) in each multiple myeloma patient (x-axis) in the trial identified using SC-expression, where the color code for the sub-clones is provided at the top. **(D)** Predicted viability of the combination at a clonal level for each patient where response status is provided at the bottom-strip of each facet. The left to right order of patients is the same as in panel A. (**E)** The predicted combination response in 28 multiple myeloma patients stratified by responder vs. non-responder status. (**F)** Receiver Operating Characteristic curve displaying the predicted combination response. The area under this curve, provided at the right bottom corner, denotes the overall stratification power in distinguishing responders vs. non-responders.

We succeeded in building predictive PERCEPTION response models for two out of four of these drugs (carfilzomib and lenalidomide). Using these models, we predicted the combination response for a given patient via the following two steps (Figure 4A-B): **<u>(1)</u>**<u>Predicting the combination response of each clone in that tumor:</u> We first predicted the combination response for each clone (one of three transcriptional clusters originally identified in (Cohen et al. 2021) based on its mean expression profile across all the cells composing it. To this end, we first predict the response for each of the two drugs in the combination separately based on their respective PERCEPTION models. Second, we then take the maximal killing by one of these two drugs as the predicted killing of the combination for that specific clone, following the Independent Drug Action (IDA, Ling et al. 2020) (**Figure 4B**). **(<u>2)</u>** <u>Second, having predicted the combination effect on each of the clones present in the tumor, we use these predictions to predict the overall patient’s response (</u>**<u>Figure 4A</u>**): This prediction is taken as the predicted response of the least responsive clone, assuming that it is likely to be selected by the treatment and dominate the overall tumor’s response. This notion is further motivated by observing that the predicted response of the most resistant clone indeed best stratifies the responders vs. non-responder patients among four different strategies that we devised and tested on this dataset using SC-expression (**Supplementary Note 2, Methods**). This strategy is then fixed and used in all other patients’ analyses shown herein.

Based on this approach, we have applied PERCEPTION to predict the response of all 28 patients in the trial to the combination treatment. **Figure 4D** shows the <u>predicted viability of the combination at a clonal level for each patient (in a layout corresponding to</u>**<u>Figure 4A</u>**). As evident, the resulting predicted response scores (1-predicted viability) are significantly higher in responders vs. non-responders **(Figure 4E)** and can successfully predict treatment response with fairly high accuracy (ROC-AUC of 0.827, **Figure 4F)**.

We compared PERCEPTION stratification performance to four different kinds of control modes (**Supplementary Notes 2 and 3**). First, we repeated the analysis using pseudo-bulk expression (mean expression over all the cells in the tumor) yielding a poor ROC-AUC of 0.56, which testifies to the marked benefit of harnessing SC data from patient tumors to predict their response. Second, we computed this response using the strategy we used for cell lines and PDCs, taking the mean viability across all single cells in a tumor sample, yielding a predictive signal with ROC-AUC of 0.64, which is considerably inferior to prediction accuracy obtained via the clonal based approach. (**Supplementary Note 2**). Third, as further controls, we built and tested three different types of random models, built by (1) Shuffling the viability labels in the cell lines, by (2) randomly selected gene signatures, and finally (3) using non-predictive models of other drugs (**Methods**). These models yielded significantly lower stratification power than that obtained by PERCEPTION (empirical P-values over 1000 instances of P=0.002, P<0.001, and P<0.001, respectively). Finally, we compared the PERCEPTION stratification performance with published state-of-the-art machine learning response models for cell lines and are trained only on bulk-expression (Tsherniak et al. 2017) and PERCEPTION models that are not tuned on SC-expression. We found that repeating the analysis with the above two models yields quite inferior performance, with AUCs of 0.62 ± 0.001 and 0.52 ± 0.001, respectively (**Extended Figure 6, Supplementary Note 2**).

### PERCEPTION predicts CDK inhibitor treatment response in a breast cancer clinical trial

Using the prediction approach described in the previous section, we next tested PERCEPTION’s ability to predict patient response in the FELINE breast cancer clinical trial with SC-expression profiles of 34 patient tumors (Griffiths et al. 2021). This clinical trial includes three treatment arms: endocrine therapy with letrozole (Arm A), an intermittent high-dose combination of letrozole and CDK inhibitor ribociclib (Arm B), and a continuous lower dose combination of the latter (Arm C). SC-expression and treatment response labels were available for 33 patients (Arm A=11 samples, Arm B= 11 and Arm C=11; **Table S7**). Patient response was determined via tumor growth measurements from mammogram, MRI, and ultrasound of the breast.

PERCEPTION was able to build a predictive response model only for ribociclib (with a Pearson R=0.26, P=1.5E-03, which is a bit lower than the threshold we used in the overall analysis), and thus we focused our analysis on the combination arms B and C that include it (**Figure 5A**). We processed the SC-expression profiles of the tumor cells as described in (Griffiths et al. 2021) and identified 36 transcriptional clusters/clones that are shared across the patients (**Extended Figure 7A-C, Supplementary Note 3, Methods**). Patient response was predicted in a precisely similar manner to that described above in the case of the multiple myeloma trial. As the number of patients is quite small we collated together and predicted the response of the patient pre-treatment samples in Arm B and C in aggregate. The resulting predicted viability of the non-responders is higher than that of the responders (Wilcoxon rank-sum test, one-sided P=0.05, **Figure 5B**), as expected. The PERCEPTION predictor successfully stratified the responders vs. non-responders with a quite high ROC-AUC of 0.776 (**Figure 5C**). As in the multiple myeloma case, PERCEPTION*’s* stratification performance is higher than three random control models, including (1) 1000 PERCEPTION models generated by shuffling the viability labels (P-value=0.039), (2) 1000 randomly generated gene signatures (P=0.043), and (3) 200 non-predictive PERCEPTION models (P=0.02). It also outperforms two bulk expression-based prediction models, including (1) published drug response models (Tsherniak et al. 2017) (AUC=0.60 ± 0.009) and (2) PERCEPTION bulk expression-based models that have not been tuned on SC-expression (AUC=0.63 ± 0.012) (**Methods, Extended Figure 7D**). Notably, computing the response using the strategy used for cell lines and PDCs that is based on computing the mean viability across all single cells yields a decent AUC of 0.735, which while being a bit lower than that achieved by the clonal response in this case, still markedly outperformed that numerous control models (**Extended Figure 7E, Supplementary Note 3**).

**Figure 5:**
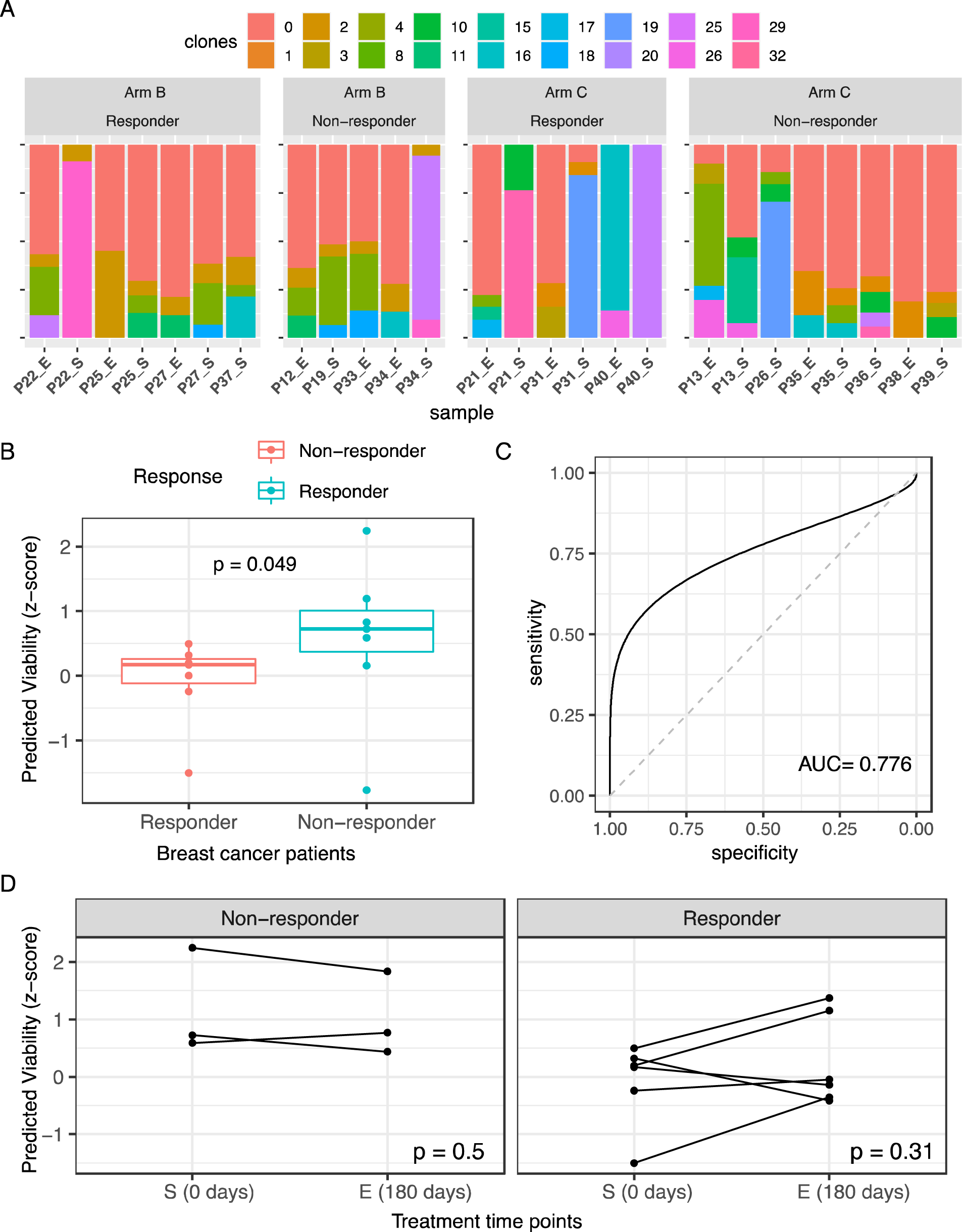
PERCEPTION stratifies responders vs. non-responders of the combination therapy arms in the FELINE clinical trial. **(A)** Distribution of abundance of malignant sub-clones (y-axis) in each breast cancer patient from the combination arms B and C (x-axis) in the trial identified using SC-expression, where the color code for the sub-clones is provided at the top. In the x-axis, the labels are a combination of the patient id and the time point at which the sample was collected (“_S” - day 0 and “_E” - day 180). (**B)** The predicted combination response in 14 breast cancer patients (samples collected at day 0), stratified by responder vs. non-responder status. (**C)** Receiver Operating curve displaying the predicted combination response. The area under this curve, provided at the right bottom corner, denotes the overall stratification power in distinguishing responders vs. non-responders. (**D**) One-to-one comparison of predicted viability (y-axis) in response to ribociclib treatment at time points S (0 days) and E(180 days) (x-axis) in non-responders (N=3) and responders (N=6) separately.

Along with the pre-treatments samples collected at the time of screening (S), the clinical trial included the SC-expression of samples collected at two post-treatment time points, on day 14 (M) and on day 180 at the end of the trial (E). Of the patients we analyzed, eight (6 responders and 3 non-responders) patients had paired SC-expression at day 0 and day 180. Of note, within the responders, there is an overall trend (non-significant, given the very small sample size), predicted viability increases from day 0 to day 180 (Wilcoxon Signed Rank test, two-sided P=0.30; Mean fold change between S and E, FC=0.35 **Figure 5D**) where in contrast, no trend was observed within non-responders (P=0.5, FC=1.17, **Figure 5D**, regression interaction P=0.11). The reduced response trend observed specifically among the responders after 180 days could possibly indicate a selection pressure for resistant tumor clones. However, we obviously note the limited samples in our cohort and thus this observation needs to be replicated in a larger cohort.

### PERCEPTION quantifies the development of resistance to multiple tyrosine kinase inhibitors trial in lung cancer patients

We next tested if PERCEPTION can capture the development of clinical resistance during targeted therapy treatment in patients. To this end, we analyzed a recently published cohort with a scRNA-seq profile of 24 lung cancer patients with 14 pre- and 25 post-treated biopsies (Maynard et al. 2020) (**Extended Figure 8A-F, Table S8**). In total, patients in this cohort were treated with four different tyrosine kinases including erlotinib (a 1^st^ generation EGFR inhibitor), dabrafenib (a BRAF inhibitor), osimertinib (3rd generation EGFR inhibitor), and trametinib (a MEK inhibitor). Based on the notion that the resistance to these target therapies usually increases as the treatment prolongs, we hypothesized that the resistance predicted in a given post-treatment biopsy would increase as elapsed treatment time (the number of days from the start of the treatment to the day biopsy was collected) increases.

We first analyzed all samples available, comparing the pre and post-treatment cohorts as a whole (as the majority of the samples were not matched). To this end, for each such post-treatment biopsy, we defined the estimated “*Extent of Resistance’’* to a given treatment as the difference between its predicted viability vs. the baseline predicted viability, where the latter is computed as the mean predicted viability across all pre-treatment biopsies. Aligned with our hypothesis, we find that the *extent of resistance* to treatment is strongly positively correlated with the elapsed treatment time, but notably, only in the patients that have been reported to acquire resistance in the original trial (Progressive Disease, Spearman Rho=0.634, P=0.026, **Figure 6A**, N=17). We also found that this positive *co*rrelation between the elapsed treatment time and the estimated extent of resistance holds true when patients receiving different drugs are analyzed separately (**Extended Figure 9A**), controlling for prior treatments (**Extended Figure 9B**), when individual patients are analyzed separately (**Extended Figure 9C**) and the stage (**Extended Figure 9D**). We also note that the extent of resistance is significantly higher in post-treatment biopsies collected from the patients with Progressive Disease vs Residual disease (Wilcoxon Rank sum P<0.002, Stratification ROC-AUC=0.88, **Figure 6B**). Notably, we do not observe this strong positive correlation but rather a negative trend in patients that have responded well to the treatment (Residual Disease, N=7, Spearman Rho= -0.67, P=0.11, **Figure 6A**). This increase in the extent of resistance to treatment with elapsed treatment time specifically in patients that acquire resistance is not significant when considering only bulk-expression from tumors (**Extended Figure 9B, Supplementary Note 4**).

**Figure 6:**
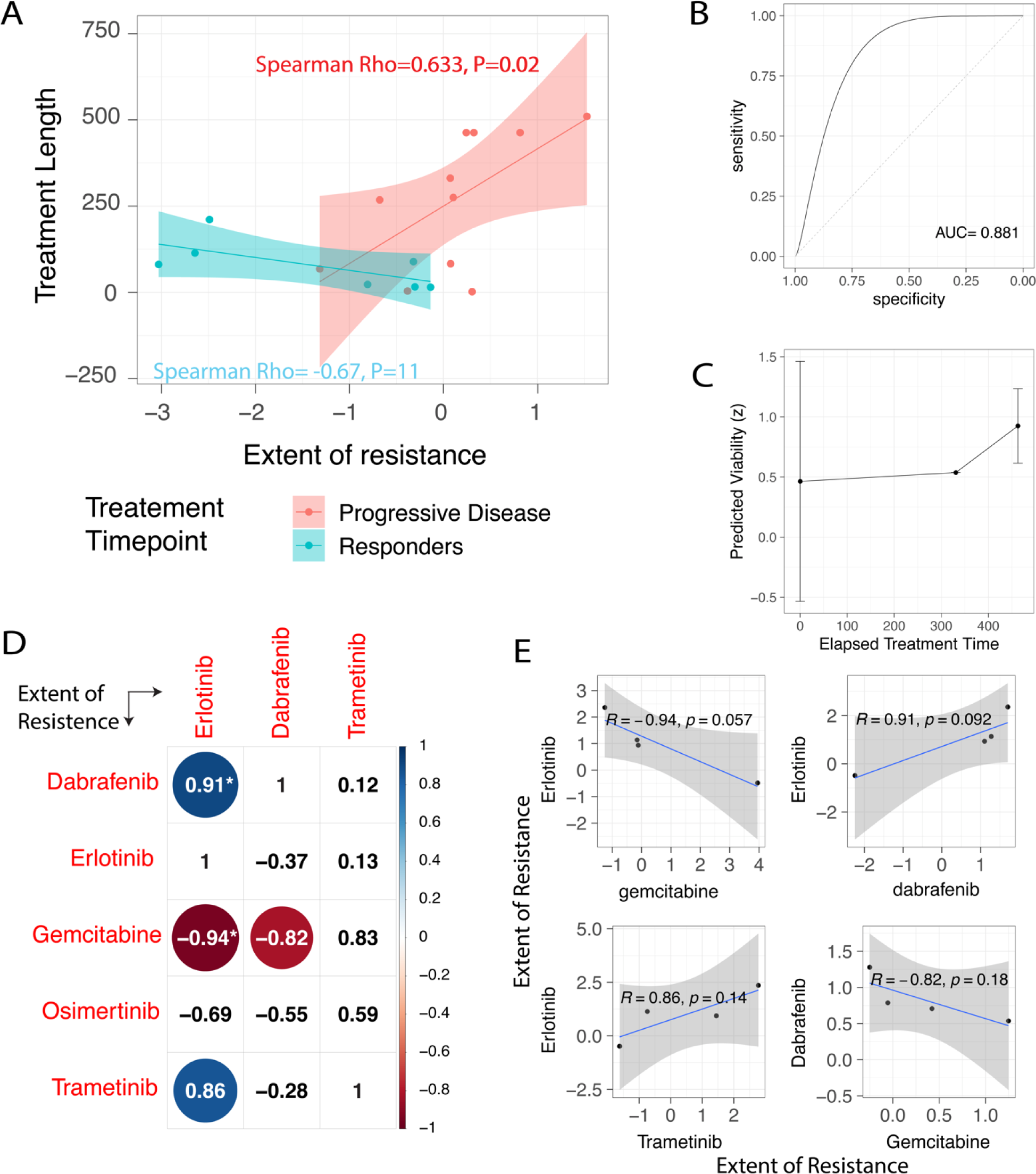
Predicting the development of resistance to tyrosine kinase inhibitors (TKIs) during treatment in NSCLC patients. **(A)** The extent of resistance to a treatment from the baseline (X-axis) is correlated with the treatment elapsed time (Number of days from the start of the treatment before the biopsy was taken) (Y-axis). The points and line colors denote whether the biopsy is from patients with progressive disease or responders. (**B)** Receiver Operating curve depicting the stratification power in distinguishing the biopsies from the patients that acquire resistance vs biopsies from the patients that responded based on their extent of resistance. (**C)** The case of Patient TH179 with multiple biopsies is presented where the predicted viability in 14 pre (day 0) and 4 post-resistant tumors at day 331 (N=1) and day 463 (N=3) to dabrafenib are shown. (D) Correlation matrix of the extent to resistance among drugs available in the trial across all the patients that have acquired resistance to this treatment. The strength of the correlation (Pearson R) is provided in the respective box and has been represented by the size of the circle, where the color represents whether the correlation coefficient is negative or positive (red and blue, respectively). We have computed this for drugs with at least three resistant patients (# of patients=4, 4, and 3, respectively). The drugs with correlations significance threshold before FDR correction <0.1 are indicated by a “*”. (**E)** Correlation plot of drug pairs with cross-resistance or cross-sensitivity with a significance threshold before FDR correction <0.2.

We next analyzed the subset of patients with matched biopsies, including five patients with two biopsies each and one patient with four biopsies. We find that the correlation pattern between treatment elapsed time and estimated extent of resistance holds in the matched cases, specifically in patients that acquire resistance (Regression Interaction P=0.003). Of particular interest is a case of a single patient (TH179), treated with dabrafenib, that had four biopsies at two different time points and developed progressive disease. The predicted viability to dabrafenib of the four tumor biopsies taken after 331 and 463 days of start of treatment is significantly higher than pre-treatment biopsies (**Figure 6C**). Furthermore, the predicted viability of all three biopsies from day 463 is significantly higher than the biopsy from day 331. Taken together, these results quantify the emergence of treatment resistance as the disease progresses and testify to the ability of PERCEPTION analysis to capture it.

To prioritize candidate drugs available in this cohort whose treatment may overcome the acquired resistance, we next asked if the development of resistance to a drug can induce either cross-sensitivity or cross-resistance to the other drugs (Plucino et al. 2012) in the cohort. To this end, we focused on the patients (**Table S8**) that eventually acquired resistance and computed the correlation among each drug’s extent of predicted resistance across these patients (**Figure 6D, Methods**). In the patients we analyzed, PERCEPTION predictions suggest that the development of resistance to erlotinib would induce a cross-sensitivity to gemcitabine (**Figure 6E**, Top-Left panel, Pearson’s R= -0.94, P=0.06) and cross-resistance to dabrafenib (**Figure 6E**, Top-Left panel, Pearson’s R=0.91, P=0.09). A literature survey (**Methods**) revealed that gemcitabine treatment can overcome erlotinib resistance in cancer cell lines through downregulation of Akt (Bartholomeusz et al. 2011). In patients, a combination of gemcitabine+erlotinib in pancreatic cancer in phase III trial has shown a higher overall and progression-free survival (Moore et al. 2016, Shin et al. 2016). In contrast, the addition of trametinib to erlotinib did not significantly improve survival in a phase I/II clinical trial (Luo et al. 2021). In sum, our analysis supports the possibility that erlotinib resistance may induce cross-sensitivity to gemcitabine, suggesting its further future testing.

## Discussion

We present a first of its kind computational framework and pipeline for predicting patient response to cancer drugs at single-cell resolution. We demonstrate its application for predicting response to monotherapy and combination treatment at the level of cell lines, patient-derived primary cells, and in predicting patient response in three recent clinical cohorts, spanning multiple myeloma, breast cancer, and lung cancer. Applying the PERCEPTION models to patients’ tumor data, we observed that incorporating the transcriptional clonal information of the tumor into the prediction process improves the overall accuracy. For a given patient, the transcriptional clone with the worst response, that is the most resistant pre-treatment clone, best represents their overall response to treatment.

We note that computing response cell lines using the strategy we used for predicting the response in clinical trials (most-resistant-clone strategy) can also quite successfully stratify resistant vs sensitive cell-lines, however, with markedly lower performance (**Extended Figure 10-11, Supplementary Note 5**) than the mean-response strategy that we used for predicting cell-lines response (mean predicted response computed over all the individual single cells from a cell line). One possible explanation for the difference in the performance of mean vs most-resistant-clone based prediction strategies observed in the cell-lines vs patient’s data could be that the clinical responses are measured at much longer time-scales in the patients (months) than in the cell lines (within days), thus providing time for the selection of the most resistant clone and making the clonal based strategy much better fit to predict patients response than the mean-based one.

The scope and power of our analysis are currently considerably limited by the scarce availability of pre-treatment SC-expression patient datasets with treatment response labels. One can quite confidently estimate that the accuracy and breadth of SC-based drug response predictors will markedly increase in the foreseen future with the growing availability of such data. In essence, it is another incarnation of the chicken and egg scenario – these relatively costly datasets will only be generated on a large scale when their clinical utility becomes more apparent, and the current paucity of these datasets yet impedes further progress. Hence, the current demonstration of their potential value, coupled with the basic intuition that one needs to target multiple clones in tumors to achieve long-enduring responses, will hopefully serve to drive the generation of relevant datasets in clinical settings moving forward. Considering that the average annual cost of treating a cancer patient in the US is currently around $150K, the current cost of about $15K for sequencing a tumor to optimize treatment is one order of magnitude smaller, and at least in our minds, a well-justified expense that is ought to be carefully studied and considered moving forward.

Consequently, one can further expect that SC-based drug response predictive models would further improve when such datasets would become more available. But beyond that, they could be further improved by considering cancer type-specific cell lines, whenever a large number of such models become available for each cancer type. We note that the quality of our response models would also depend on the quality of the SC-expression profiles available e.g., their depth, drop-out rates, etc. Of note, we deliberately have chosen not to impute the SC data given the recent reports that dropouts are limited to non-UMI-based SC-expression methods and otherwise likely reflect true biological variation (Svensson et al. 2020, Cao et al. 2021). A key limitation of our pipeline is a lack of ability to predict drug effects on immune and normal cells in the tumor microenvironment, which is obviously needed to estimate the toxicity and side effects of different combinations. A major push to future SC-based precision oncology development will come from large-scale drug screens of drugs in noncancerous cell lines, currently very scarcely available, which will then enable the construction of predictors of drug effects on non-tumor cells, using an analogous pipeline to the one presented here for tumor cells. Finally, our results demonstrate that tracking the drug response expression in post-treatment biopsies could help follow the evolution of drug resistance at a single-cell resolution and help guide the design of future personalized combination treatments that could significantly diminish the likelihood of resistance emergence.

In summary, this study is the first to demonstrate that the high resolution of information from scRNA-seq could indeed be harnessed to build drug response models that can predict effective targeted therapies for individual cancer patients in a data-driven manner.

## Methods

### Data collection

We first collected the bulk-expression and drug response profiles generated in cancer cell lines curated in the DepMap (Tsherniak et al. 2017) consortium from Broad Institute (version 20Q1, https://depmap.org/portal/download/). The drug response is measured via area under the viability curve (AUC) across eight dosages and measures via a sequencing technique called PRISM (Corsello et al. 2020). In total, we mined 488 cancer cell lines with both bulk-transcriptomics and drug response profiles. We next mined SC-expression of 205 cancer cell lines (280 cells per cell line) generated in (Kinker et al. 2020) and distributed via the Broad Single-cell Portal. The metadata, identification, and clustering information were also mined from the same portal (https://singlecell.broadinstitute.org/single_cell/study/SCP542/pan-cancer-cell-line-heterogeneity#study-download).

### The PERCEPTION pipeline

A response model for a drug is built via the following two steps: **1. Learn from bulk** and **2. optimize using SC expression**. We first divided all the cancer cell lines into two sets - 1. Cell lines where bulk-expression is available, and SC-expression is not available (N=318) 2. Cell lines where SC-expression is available (N=170). The first set is used during learning from bulk (Step 1, expanded below) and the second in optimizing using SC expression (Step 2).

**Step 1: Learn from bulk:** As a feature selection step, we first identified genes whose bulk-expression is correlated with drug viability profile (using the Pearson correlation). We considered the Pearson correlation *Pc(d, g)* between drug *d* and gene *g* as a measure of information in a gene expression profile and ranked each gene based on the strength of the correlation. While considering the top X genes, where X is a hyperparameter optimized in the next step, we built a linear regression model regularized using elastic net to predict the response to *d* in five-fold cross-validation, as implemented in R’s glmnet (Friedman et al. 2010).

**Step 2: Optimize using SC-expression:** We built the above model using a Bayesian-like grid search of various possible values for X (range 10-500), where the model with the best performance using SC-expression input of 169 cell lines (left one out for testing) was chosen. We finally performed measured the model performance in a leave one our cross-validation using the left-out cell line, which was not used in either model building or hyperparameter optimization. Performance was measured using Pearson’s correlation between the predicted response and the actual response.

### Cross-platform comparison of PERCEPTION performance

The pharmacological drug screens performed by the PRISM and GDSC studies are based on two independent platforms. The GDSC data were downloaded from the DepMap portal (Downloaded: April 15, 2020, https://depmap.org/portal/download/). To compare the performance of PERCEPTION across two independent screening platforms and test if the expression signature captured by our drug response models can be translated across the domains, we tested according to the following multi-step procedure:

1. Of the 347 cell lines in common with drug response in both GDSC and PRISM, there are 120 cell lines with SC-expression data in (Kinker et al. 2020). We selected at random 80 cancer cell lines with SC-expression data and pharmacological screens in GDSC and PRISM,
2. We considered all the drugs (N=191) that were screened in both PRISM and GDSC, from which we selected a subset of drugs (N=28) with a concordant response between PRISM and GDSC (Pearson rho > 0.3 and p-value < 0.05; at least 20 cell lines with responses per drug in both GDSC and PRISM) in the 267 cell lines in common between the two screens excluding the cell lines in the testing set.
3. For each of the drugs selected in step 2, we ran the PERCEPTION pipeline with one necessary change in the set of cell lines used. Specifically, in Step 2, the parameters were optimized on SC-expression of 90 cell lines (excluding the 80 test cell lines) instead of the default 170 cell lines which have response data in PRISM.
4. Finally, we applied the resulting response models to the testing dataset and compared the predicted AUC values to the experimental responses from GDSC and PRISM. We used the Pearson correlation coefficient as the measure to compare the performance between the screens and predicted responses.

### PERCEPTION prediction of monotherapy and combination response in a lung cancer cell lines screen

We first performed a qualitative test of the drug screen mined from (Nair et al. 2021), where the response is measured via the AUC of the dosage-viability curve across eight dosages. To this end, we compared this screen to a previous high-quality screen from Broad and Sanger Institute, PRISM (Corsello et al. 2020). Specifically, we leveraged the fact that the two screens have common drugs and share some cell lines. Focusing on this set of cell lines and drugs, for each drug, we computed a correlation between the viability profile in the screen from Nair et al. and PRISM (**Extended Figure 3A**). We reasoned that the drugs with correlated profiles in the two screens (Pearson Rho>0.3, defined as concordance score) are consistent across the two screens, suggesting that they are high quality. Independently, we also note that the concordance score of drugs’ response profile across screens is correlated with our predictive performance (Pearson Rho>0.39; P<0.019, **Extended Figure 2A**), suggesting that our model is capturing the robust signal across screens of these drugs. In this screen, we defined AUC data points <1 as high-confidence and we filtered out the other data points as typically an AUC larger than 1 is due to noise in the data as we see higher variability in doses that do not inhibit.

We focused on data points for 14 FDA-approved drugs in 21 cell lines that passed the above filter for which we could build predictive PERCEPTION models (Pearson’s R>0.3, P<0.05). We assessed their predictive performance vs. drug screen data measured for monotherapy and two-drug combinations of these drugs across 21 lung cancer cell lines in five dosages (**Table S5**). Using SC-expression for these lung cancer cell lines profiles in (Kinker et al. 2020) (300 cells per cell line), we used the PERCEPTION models to predict the response to each drug in each cell line by computing the mean predicted viability across all the single cells of that cell line. We next tested PERCEPTION’s ability to predict the response to combinations of these 14 drugs studied in this screen (**Table S5**). A combination response in a given cell line was predicted by adopting the independent drug action (IDA) model across all the single cells from that cell line (Ling et al. 2020); i.e., the predicted combination response of N drugs is the effect of the single most effective drug in the combination. Performance was measured using ROC-AUC. Throughout our work, combination response is predicted using the IDA principle.

### PERCEPTION’s prediction in patient-derived head and neck cancer cell lines

The single-cell expression of the five head and neck squamous cell cancer (HNSC) patient-derived cell lines and their treatment response for eight drugs and combination therapy at two different dosages were obtained from (Suphavilai et al. 2020). For these drugs, PERCEPTION was unable to build drug response models with Spearman correlation between their predicted vs. experimental viability greater than 0.3 using PRISM screens. Therefore, we incorporated two main changes to the PERCEPTION pipeline:

1. Drug response from GDSC screens (response from > ∼800 cell lines for these drugs) were used to build models,
2. Only the top 3000 highly expressed genes (with fewer dropouts in HNSC dataset) in common between the bulk expression and PDC datasets were considered in the pipeline. For the drugs for which PERCEPTION was able to build models, we applied the models on the PDC cell lines and obtained the predictions for each individual cell. The patient-level monotherapy response for a given drug is represented by the mean response of all the cells included in a patient’s PDC sample. In the case of drug combinations, for a given cell, its combination response is computed using IDA, i.e., the predicted combination response of N drugs is the effect of the single most effective drug in the combination (Ling et al. 2020, IDA). The patient-level combination response was represented by the mean of the combined response of all the cells in a patient’s PDC sample.

### Predicting combinations response in multiple myeloma patients

Response labels, SC-expression of patients’ tumors (MARS-seq), clustering annotation and mean cluster expression were mined from the original publication (Cohen et al. 2021). We only used and focused on the cells annotated as malignant. The steps to predict the combination response of a patient can be divided into a two-step process: Step 1. *Predict the combination response of each clone in that tumor*, Step 2. *Predict patient’s response from the clone-level combination response*. To this end, we first tried to build PERCEPTION response models for the four treatments used in the combination therapy. However, we were able to build predictive response models for only carfilzomib and lenalidomide. We first predicted the combination response for each transcriptional cluster (or simply referred to here as a “clone”). To this end, we predicted the response for each of the two drugs separately and computed the killing using the Independent Drug Action (IDA) principle i.e., the predicted combination response of N drugs is simply the effect of the single most effective drug in the combination (Ling et al. 2020). To overcome the challenge of the discrepancy of dosage used in the clinic vs. pre-clinical testing where our models are built, we z-scale our predicted response profile of a drug across clone, where this z-score predicted response represents the relative response of a clone compared to all the other available in the cohort.

In Step 2, we use this clone-level combination killing profile in a patient to predict the overall patient’s response. We considered the predicted response of the least responsive clone found in each patient as that overall patient’s response. This is based on the notion that it would be selected by the treatment and dominate the overall tumor. Performance was measured using ROC-AUC. For our model building control, we built random models using either shuffled labels, randomized features in the regression model, or an unpredictive model of another drug in the screen for 1000 times and computed the number of times that the stratification power denoted by AUC is higher than our original model. This proportion is provided as an empirical P-value.

### Predicting combinations response in breast cancer clinical trial analysis

The pre-filtered 10X based single-cell RNAseq count data and the cell type annotations of the 65 breast cancer samples (34 patients) were downloaded from GEO (GSE158724). We considered only the cells annotated as tumor cells in our analysis. As defined in the primary publication of the dataset (Griffiths et al. 2021), we applied Seurat (v.4.0.5). We filtered out samples with fewer than 100 cells. We used the reciprocal principal-component analysis integration workflow to integrate the tumor cells from the remaining samples (Hao et al. 2021). The data were normalized using the SCTransform function and the top 5000 variably expressed genes and the first 50 PCs were used in the anchor-based integration step. The first fifty PCs and a k.param value of 20 were used to identify neighbors and the resolution was set to 0.8 to find distinct clusters. We identified 36 different clones, of which only 16 clones were found in the pretreated samples from patients in Arms B and C. The SC-expression of 16 clones was considered in the drug response prediction analysis. The patient response information was obtained from Table S12 in (Griffiths et al. 2021).

The default PERCEPTION pipeline was used to build drug response models except for a single change. The top ∼2500 highly expressed genes (ranked by total number of non-zeroes across all the cancer cells) in the breast cancer dataset that are in common with the cancer cell line bulk expression data were used in the pipeline. The resulting models were used to predict response at the patient level in a similar manner to what we did for the multiple myeloma data. The controls for model building were also tested for the breast cancer data similarly to the testing we did for the multiple myeloma data.

### Building bulk-based drug response models to distinguish responders from non-responders

We built bulk-based drug response models to compare their performance vs PERCEPTION models in stratifying responders from non-responders in the two clinical trials. To build drug response models based on bulk expression data, we considered all ∼500 cell lines with bulk expression and PRISM-based drug response. For each drug, we randomly divided the data in training (1/3rd of the cell lines) and test set (2/3rd of the cell lines). As a feature selection step, we first identified genes whose bulk-expression is correlated with drug viability profile (Pearson R) in the training set. We considered *Pc(d, g)* as a measure of information in a gene expression profile and ranked each gene based on the strength of the correlation. While considering the top 100 genes, we built a linear regression model regularized using elastic net to predict the response to in leave one out cross-validation, as implemented in R’s glmnet (Friedman et al. 2010). The resulting model performance was validated on the testing dataset.

To build state-of-the-art drug response models as defined in (Tsherniak et al. 2017), we generated random-forest-based models in a similar framework as defined above. To make sure that the gene features used in the resulting model predictors are actually detected to be expressed in the patient SC-dataset, we consider genes that overlap in both the cell line bulk expression data and patient SC-dataset to build the models. For each drug, we repeated the above model-building steps 100 times and presented the mean and standard error of their performances in stratifying responders from non-responders in their respective clinical trials.

### Predicting the development of resistance to multiple tyrosine kinase inhibitors trial in lung cancer patients

The SC-expression profiles of 39 biopsies from 25 patients were provided by the authors of (Maynard et al. 2020). The clinical annotations were mined from the original publication, specifically Supplementary Table 2. Similar to previous sections, we focused only on the subset of single cells labeled in the publication as malignant. Seurat clustering was performed with the resolution = 0.8, dims = 10, number of features = 2000, scale.factor = 10000, log normalization method with minimum cells in a cluster required to be > 3 and minimum features required to be > 200, to identify a total of 16 clones. The expression of each transcriptional cluster/clone in a patient is the averaged expression across all the single cells associated with that cluster in that given patient. We successfully built drug response models for dabrafenib, erlotinib, gemcitabine, osimertinib and trametinib. The response observed in the most resistant clone of a patient is considered as the PERCEPTION’s predicted response. We primarily studied the development of drug resistance in the trial. To this end, we defined a term called “Extent of Resistance” of a drug, which is a difference between a drug’s predicted viability from PERCEPTION and the predicted baseline viability. The predicted baseline viability is defined as the average predicted viability of the respective treatments in all treatment-naive samples. This difference in response from the naive state denotes the extent of resistance and is thus named accordingly. We computed correlations using both Spearman and Pearson to test and identify robust correlations.

### Literature survey of cross-resistance and cross-sensitivity

To search for evidence available in published papers for a cross-resistant or cross-sensitive drug pair, we used the search term “drug X AND drug Y” e.g., erlotinib AND gemcitabine, in the PubMed search portal https://pubmed.ncbi.nlm.nih.gov/ on December 26, 2021. The resulting clinical trials in the first fifty matches, sorted by best match, were manually surveyed for outcomes. For pre-clinical evidence for or against, non-clinical studies testing the combinations were manually surveyed.

## Data availability

The entire collection of the processed datasets used in this manuscript, including pre-clinical models of cancer cell lines and PDCs, can be accessed via a Zenodo repository that will be provided upon request or upon publication.

## Code and data availability

The study’s scripts to replicate each step of results and plots will be provided upon publication. We used open-source R version 4.0 to generate the figures. Wherever required, commercially available Adobe Illustrator 23.0.3 (2019) was used to create the figure grids.

## Acknowledgments

This research was supported in part by the Intramural Research Program of the National Institutes of Health, NCI, NIH grants R01CA231300 (T.G.B.), R01CA204302 (T.G.B.), R01CA211052 (T.G.B.), R01CA169338 (T.G.B.), and U54CA224081 (T.G.B.). This work used the computational resources of the NIH HPC Biowulf cluster (http://hpc.nih.gov). We acknowledge and thank the National Cancer Institute for providing financial and infrastructural support. Thanks to Kun Wang, Sheila Rajagopal and Ze’ev Ronai for their valuable feedback and discussion. Thanks to Jason I. Griffiths and Andrea H. Bild for clarifying the patient response data in (Griffiths et al. 2021) and for their helpful feedback.

## Conflict of interest

E.R. is a co-founder of Medaware, Metabomed, and Pangea Therapeutics (divested from the latter). E.R. serves as a non-paid scientific consultant to Pangea Therapeutics, a company developing a precision oncology SL-based multi-omics approach. T.G.B. is an advisor to Array/Pfizer, Revolution Medicines, Springworks, Jazz Pharmaceuticals, Relay Therapeutics, Rain Therapeutics, Engine Biosciences, and receives research funding from Novartis, Strategia, Kinnate, and Revolution Medicines. The rest of the authors declare no conflict of interest.

## References

Adam G, Rampá šek L, Safikhani Z et al. Machine learning approaches to drug response prediction: Challenges and recent progress. NPJ Precision Oncology 2020; 4: 19.

Arya AK, El-Fert A, Devling T, et al. Nutlin-3, the small-molecule inhibitor of MDM2, promotes senescence and radiosensitises laryngeal carcinoma cells harbouring wild-type p53. British Journal of Cancer 2010; 103(2) : 186–195.

Bartholomeusz C, Yamasaki F, Saso H, et al. Gemcitabine overcomes erlotinib resistance in EGFR-overexpressing cancer cells through downregulation of Akt. Journal of Cancer 2011; 2: 435.

Bhinder B, Gilvary C, Madhukar NS, Elemento O. Artificial intelligence in cancer research and precision medicine. Cancer Discovery 2021; 11(4):900–915.

Cao Y, Kitanovski S, Küppers R, Hoffmann D. UMI or not UMI, that is the question for scRNA-seq zero-inflation. Nature Biotechnology 2021; 39(2): 158–159.

Castro LNG, Tirosh I, Suvà ML. Decoding cancer biology one cell at a time. Cancer Discovery 2021; 11(4): 960–970.

Cheng C, Zhao Y Schaafsma, et al. An EGFR signature predicts cell line and patient sensitivity to multiple tyrosine kinase inhibitors. International Journal of Cancer 2020; 147(9):2621–2633.

Cohen YC, Zada M, Wang SY, et al. Identification of resistance pathways and therapeutic targets in relapsed multiple myeloma patients through single-cell sequencing. Nature Medicine 2021; 27(3): 491–503.

Corsello SM, Nagari RT, Spangler RD, et al. Discovering the anticancer potential of non-oncology drugs by systematic viability profiling. Nature Cancer 2020; 1(2); 235-248.

de Witte, CJ, Valle-Inclan JE, Hami N, et al. Patient-derived ovarian cancer organoids mimic clinical response and exhibit heterogeneous inter-and intrapatient drug responses. Cell Reports 2020; 31(11):107762.

Friedman J, Hastie T, Tibshirani R. Regularization paths for generalized linear models via coordinate descent. Journal of Statistical Software, 2010; 33(1), 1.

Fustero-Torre C, Jiménez-Santos MJ, García-Martín S, et al. Beyondcell: targeting cancer therapeutic heterogeneity in single-cell RNA-seq data. Genome Medicine 2021; 13:187.

Garnett MJ, Edelman EJ, Heidorn SJ, et al. Systematic identification of genomic markers of drug sensitivity in cancer cells. Nature 2012; 483(7391): 570–575.

Ghandi M, Huang FW, Jané-Valbuena J, et al. Next-generation characterization of the Cancer Cell Line Encyclopedia. Nature 2019; 569(7757): 503–508.

Griffiths JI, Chen J, Cosgrove PA, et al. Serial single-cell genomics reveals convergent subclonal evolution of resistance as patients with early-stage breast cancer progress on endocrine plus CDK4/6 therapy. Nature Cancer 2021; 2(6): 658–671.

Haas L, Elewaut A, Gerard CL, et al. Acquired resistance to anti-MAPK targeted therapy confers an immune-evasive tumor microenvironment and cross-resistance to immunotherapy in melanoma. Nature Cancer 2021; 2(7): 693–708.

Hao Y, Hao S, Andersen-Nissen, et al. Integrated analysis of multimodal single-cell data. Cell 2021; 184(13):3573–3587.

Heitzer E, Haque IS, Roberts CES, Speicher MR. Current and future perspectives of liquid biopsies in genomics-driven oncology. Nature Reviews. Genetics 2019; 20(2):71–88.

Huang K, Xiao C, Glass LM, Critchlow CM,. Machine learning applications for therapeutic tasks with genomics data. Patterns 2021; 2(10):100328.

Kim KT, Lee HW, Lee HO, et al. Application of single-cell RNA sequencing in optimizing a combinatorial therapeutic strategy in metastatic renal cell carcinoma. Genome Biology 2016; 17:80.

Kinker GS, Greenwald AC, Tal R et al. Pan-cancer single-cell RNA-seq identifies recurring programs of cellular heterogeneity. Nature Genetics 2020 52(11);1208–1218.

Ledergor G, Weiner A, Zada M, et al. Single cell dissection of plasma cell heterogeneity in symptomatic and asymptomatic myeloma. Nature Medicine 2018; 24(12): 1867–1876.

Ling A, Huang RS. Computationally predicting clinical drug combination efficacy with cancer cell line screens and independent drug action. Nature Communications 2020; 11(1): 5848.

Luo J, Makhnin A, Tobi Y, Ahn L, et al. Erlotinib and trametinib in patients with EGFR-mutant lung adenocarcinoma and acquired resistance to a prior tyrosine kinase inhibitor. JCO Precision Oncology 2021; 5: 55–64.

Maynard A, McCoach CE, Rotow JK, et al. Therapy-induced evolution of human lung cancer revealed by single-cell RNA sequencing. Cell 2020; 182(5):1232–1251.

Moore MJ, Goldstein D, Hamm J, et al. Erlotinib plus gemcitabine compared with gemcitabine alone in patients with advanced pancreatic cancer: a phase III trial of the National Cancer Institute of Canada Clinical Trials Group. Journal of Clinical Oncology 2007; 25(15): 1960–1966.

Nair NU, Greninger P, Friedman A, Amzallag A, et al. A landscape of synergistic drug combinations in non-small-cell lung cancer. bioRxiv. 2021 [cited 2022 Jan 6]. p. 2021.06.03.447011. Available from: https://www.biorxiv.org/content/10.1101/2021.06.03.447011v1.abstract

Pluchino KM, Hall MD, Goldsborough AS, et al. Collateral sensitivity as a strategy against cancer multidrug resistance. Drug Resistance Updates 2012; 15(1-2): 98-105.

Shalek AK, Benson M. Single-cell analyses to tailor treatments. Science Translational Medicine 2017; 9(408): eaan4730.

Shin S, Park CM, Kwon H, Lee K-H. Erlotinib plus gemcitabine versus gemcitabine for pancreatic cancer: real-world analysis of Korean national database. BMC Cancer 2016; 16: 443.

Sade-Feldman M, Yizhak K, Bjorgaard SL, et al. Defining T cell states associated with response to checkpoint immunotherapy in melanoma. Cell 2018; 175(4): 998–1013.

Sawabata N. Circulating tumor cells: From the laboratory to the cancer clinic. Cancers 2020; 12(10):3065.

Senft D, Leiserson MDM, Ruppin E, Ronai Z. Precision oncology: The road ahead. Trends in Molecular Medicine 2017; 23(10):874–898.

Singla N, Singla S. Harnessing big data with machine learning in precision oncology. Kidney Cancer Journal 2020; 18(3):83–84.

Siravegna G, Marsoni S, Siena S, Bardelli A. Integrating liquid biopsies into the management of cancer. Nature Reviews. Clinical Oncology 2017; 14(9):531–548.

Suphavilai C, Chia S, Sharma A, et al. Predicting heterogeneity in clone-specific therapeutic vulnerabilities using single-cell transcriptomic signatures. 2020; bioRxiv https://www.biorxiv.org/content/10.1101/2020.11.23.389676v1?rss=1

Svensson V. Droplet scRNA-seq is not zero-inflated. Nature Biotechnology 2020 38(2): 147–150.

Tian L, Tomei S, Schreuder J, et al. Clonal multi-omics reveals Bcor as a negative regulator of emergency dendritic cell development. Immunity 2021; 54(6):1338–1351.

Tsherniak A, Vazquez F, Montgomery PG, et al. Defining a cancer dependency map. Cell 2017; 170(3): 564–576.

Tsimberidou AM, Fountzilas E, Nikanjam M, Kurzrock R. Review of precision cancer medicine: Evolution of the treatment paradigm. Cancer Treatment Reviews 2020a; 86:102019.

Tsimberidou AM, Fountzilas E, Bleris L, Kurzrock R. Transcriptomics and solid tumors: The next frontier in precision cancer medicine. Seminars in Cancer Biology, 2020b, in press, doi:10.1016/j.semcancer.2020.09.007.

Wensink, GE, Elias SG, Mullenders J, et al. Patient-derived organoids as a predictive biomarker for treatment response in cancer patients. NPJ Precision Oncology 2021; 5:30.

Yao Y, Xu X, Yang L, et al. Patient-derived organoids predict chemoradiation responses of locally advanced rectal cancer. Cell Stem Cell 2020 26(1): 17–26.

Zhu S, Qing T, Zheng Y, et al. Advances in single-cell RNA sequencing and its applications in cancer research. Oncotarget 2017; 8(32):53763–53779.

