## Supplementary Figures and Text for "Predicting patient treatment response and resistance via single-cell transcriptomics of their tumors"

**Supplementary Figures and Text.** Following below are eleven sets of Extended Figures and five Supplementary Notes. Any references cited can be found in the accompanying main document.

### Extended Figures

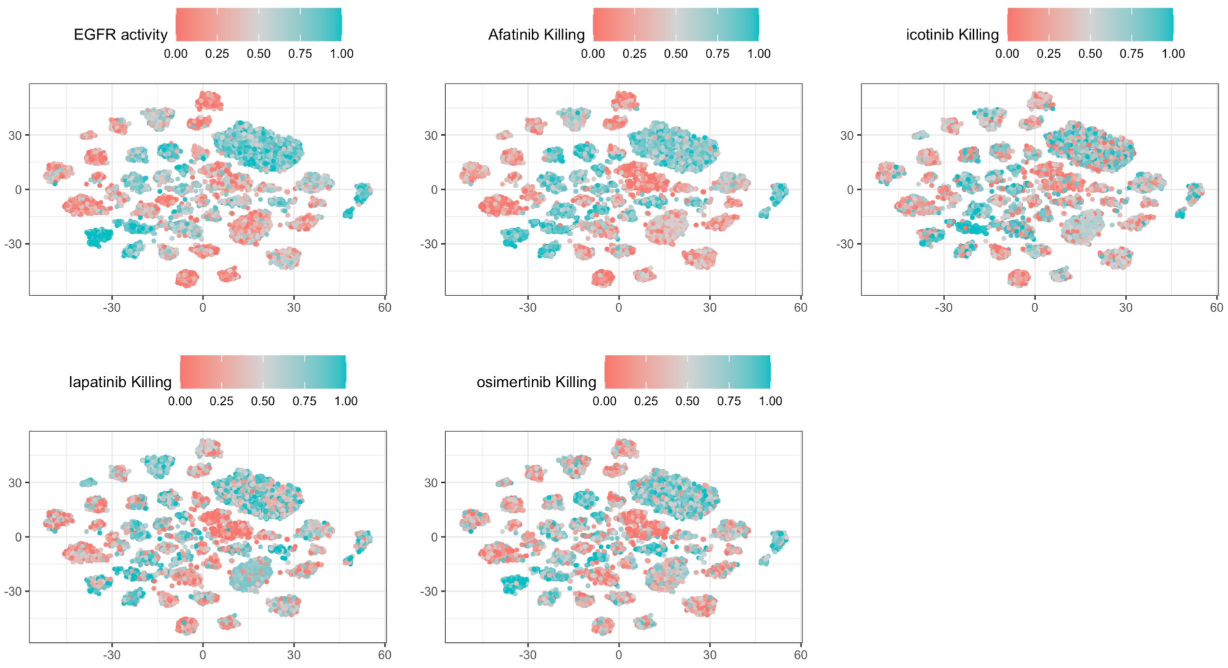


**Extended Figure 1: Visualization of PERCEPTION's ability to predict viability at four recent EGFR inhibitor vs the EGFR pathway activity at single-cell resolution:** *The top-leftmost panel visualizes the EGFR pathway activity and the other four panels visualize killing by four EGFR inhibitors, afatinib, icotinib, lapatinib, osimertinib, in every single cell (each point) via a tSNE plot, respectively. According to the legend of colors, the color of each point denotes the extent of predicted killing in the panels denoting response and the EGFR pathway activity in the top-left panel. In this figure, we provide data on 12482 individual lung cancer cells. The tSNE clustering is performed using the expression profile of all the genes.*

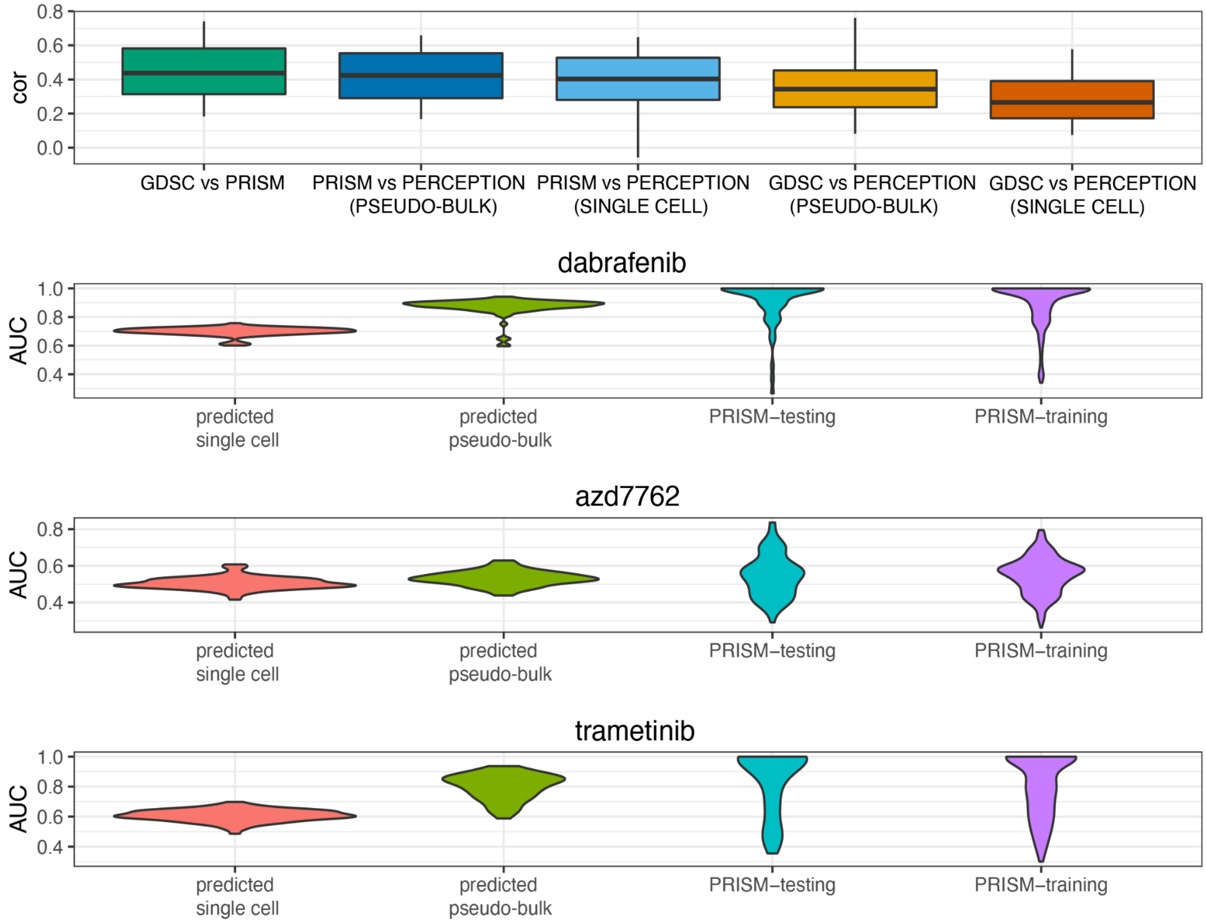


**Extended Figure 2 : PERCEPTION's performance in PRISM screens.** *A. This is an extension to Figure 2A, where we present the correlation measure via Pearson Rho (y-axis) comparing "GDSC vs PRISM", "PRISM vs PERCEPTION" and "GDSC vs PERCEPTION". Here, we also show the performance when the predictions were made at a single-cell level and pseudo-bulk (mean of single cell expression) level. Pseudo-bulk have a better performance because the random dropouts in individual cells are replaced by mean expression across cells. B-D. For the drugs dabrafenib, AZD-7762, and trametinib, we compare the distribution of predicted AUC values at single cell level (N=80, testing pan-cancer cell lines used to compare performance in GDSC), predicted AUC values at pseudo-bulk level (N=80), experimental AUC values of testing cell lines in PRISM (N=80) and experimental AUC values of training cell lines in PRISM (N=300, pan-cancer cell lines used in Step 1 of the PERCEPTION pipeline).*

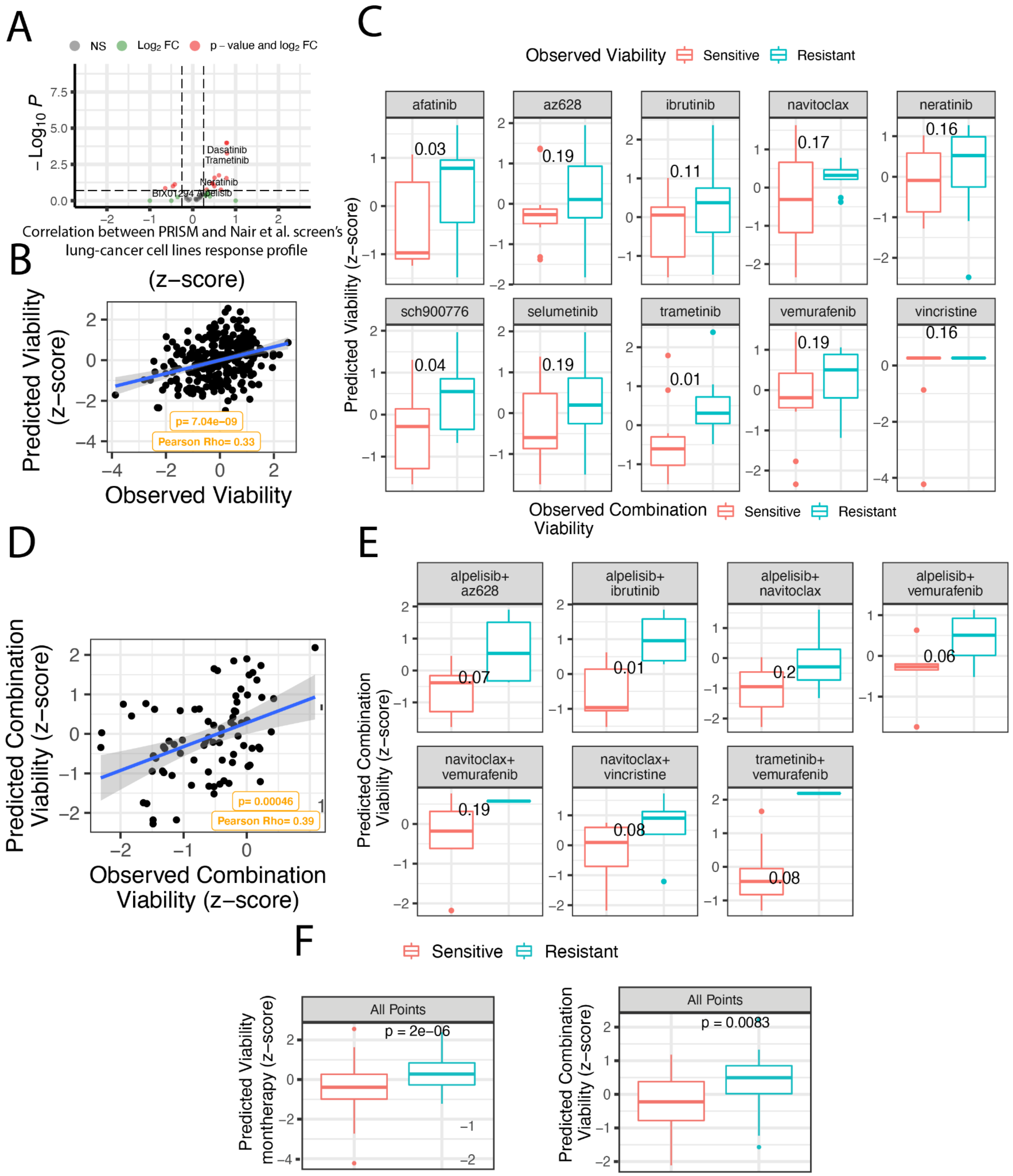


**Extended Figure 3: Screen quality control and predicted monotherapy and combination response in resistant vs sensitive lung cancer cell lines predicted by PERCEPTION for 10 drugs.** *(A) Concordance between our lung cancer screen and PRISM where the correlation Rho is provided on the x-axis and significance on the y-axis. We note that only a subset of drugs*

*shows high concordance (significantly positive correlation) between the two screens and thus we only focused on those cell lines. (**B**) Change in predicted (y-axis) vs observed (x-axis) viability, where both are centered and scaled. The Pearson correlation between predicted vs observed viability and associated significance is provided at the bottom in yellow. The best fit line is provided in blue with a 95% confidence interval in grey. (**C**) The predicted viability (x-axis) for top vs bottom 50% cell lines is defined as resistant vs sensitive cell lines, separately for each drug. A respective significance, computed using the one-tailed Wilcoxon rank-sum test is provided for each drug. PERCEPTION predicts combination response in 21 lung cancer cell lines. (**D**) Change in predicted (y-axis) vs observed (x-axis) combination viability, where both are centered and scaled. The Pearson correlation between predicted vs observed viability and associated significance is provided at the bottom in yellow. The best fit line is provided in blue with 95% confidence interval in gray. (**E**) The predicted combination viability (x-axis, scaled and centered) for top vs bottom 50 % cell lines defined as resistant vs sensitive cell lines based on observed viability, separately for 7 pairs of drug combinations. The statistical significance, computed using the one-tailed Wilcoxon rank-sum test, is provided for each drug. **(F)** The data from the distinct drugs in panel E are combined into one analysis for monotherapies and one analysis for combinations.*

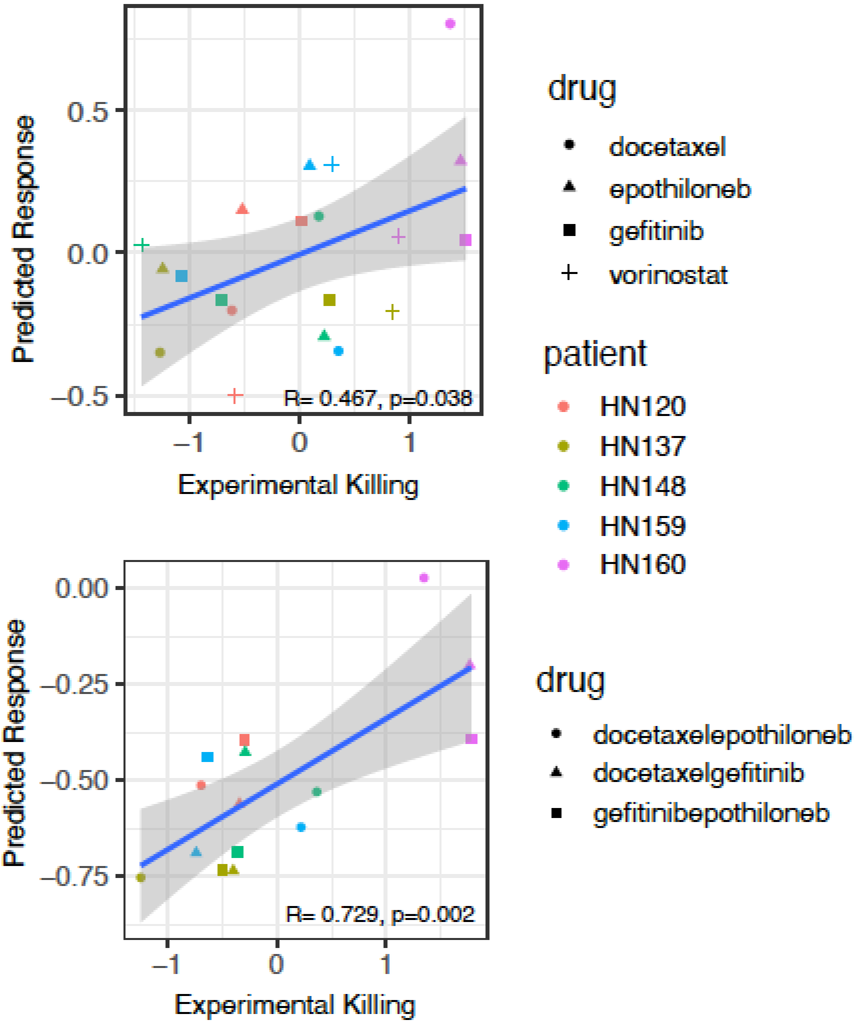


**Extended Figure 4: The predicted viability is correlated with the observed viability (median IC50) in monotherapies and combinations.** *Each scatter plot compares all the experimentally observed cell viability (x-axis; scaled for each drug treatment) to the predicted viability (y-axis; rescaled AUC value). Each dot represents the response of patient-derived cell lines (color coded) for the drugs (coded by shape) they were screened with. The Pearson correlation (R) is provided on the right side corner of each plot.*

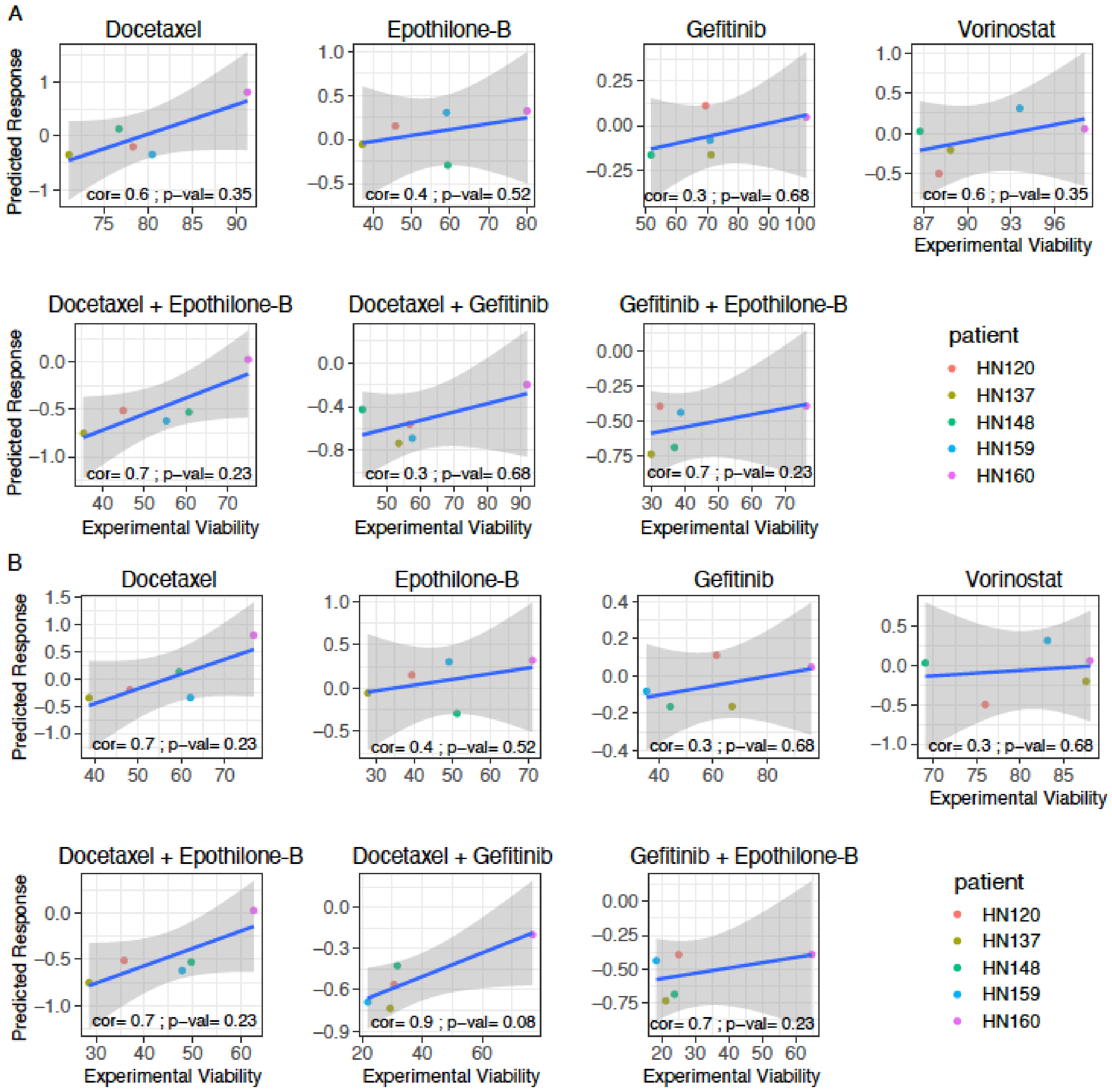


**Extended Figure 5: The predicted vs. experimental correlations obtained for individual treatments.** *Each scatter plot compares the experimentally observed cell viability (x-axis; at median IC50 concentration) to the predicted viability (y-axis; rescaled AUC value) for the four drugs docetaxel, epothilone-b, gefitinib, and vorinostat (top panel) and their combinations (Bottom Panel). Each dot represents the response of patient-derived cell lines (color coded) for the drugs (coded by shape) they were screened with. The Spearman rank correlation (rho) is*

*provided on the right side corner of each plot. This plot is provided for the following treatment - **(A)** IC50 **(B)** one-third of IC50.*

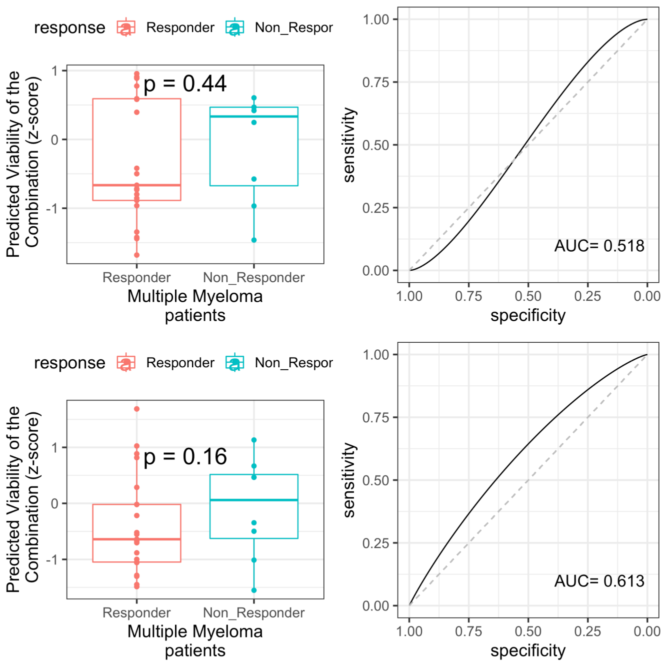


**Extended Figure 6:** *Stratification power of PERCEPTION vs published state-of-the-art machine learning response models and are trained only on bulk-expression (Tsherniak et al. 2017) and PERCEPTION models that are not tuned on SC-expression* <u>*in the multiple myeloma cohort*</u>*.*

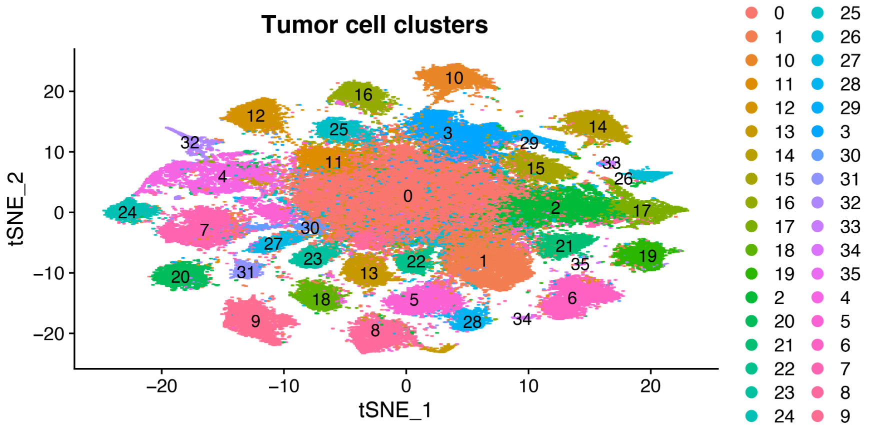


**Extended Figure 7A. The tSNE plot showing the 36 transcriptional tumor clusters identified in the breast cancer clinical trial.** *The tumor cells from 34 patients taken from three different time points were integrated and clustered using the Seurat package (Methods). The resulting 36 clusters are presented in a tSNE plot. The transcriptional clusters are color-coded and labelled as defined in the legend.*

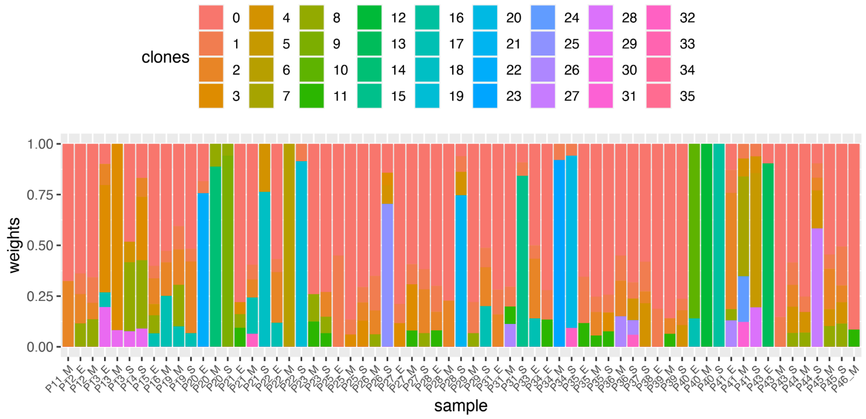


**Extended Figure 7B. Abundance of malignant sub-clones in different breast cancer samples.** *Distribution of abundance of malignant sub-clones (y-axis) in breast cancer samples (x-axis) based on SC-expression. The color code representing different sub-clones is provided in the legend (top). On the x-axis, the labels are a combination of the patient id and the time point at which the sample was collected ("_S" - day 0, "_M" - day 14 and "_E" - day 180).*

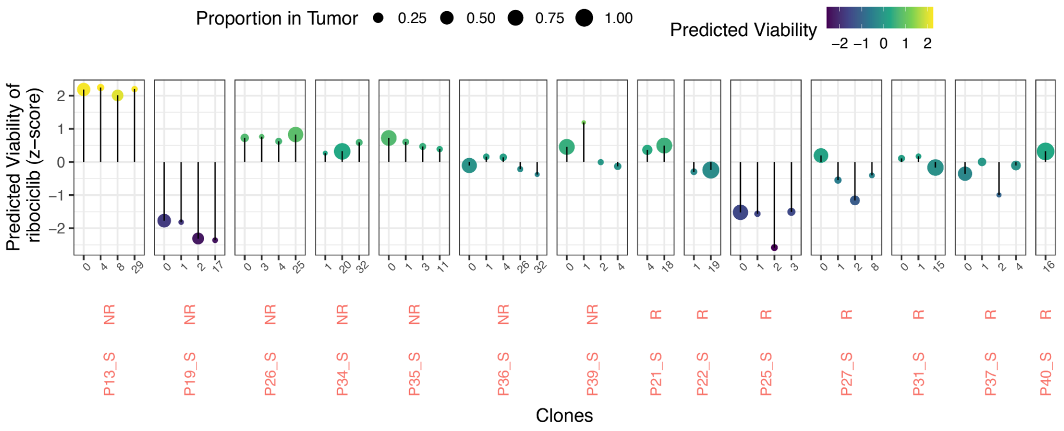


**Extended Figure 7C. Clone level response of breast cancer pre-treatment samples (day 0) in Arms B and C.** *The y-axis represents the predicted viability of ribociclib and the x-axis represents the different clones found in each patient pre-treatment samples. The response status is provided at the top strip of each column and sample name below it. The size of the dot represents the proportion of a cluster/clone compared to all the clones present in the patient. The color scale from blue to yellow represents the predicted viability, dark blue represents low viability and yellow represents high viability.*

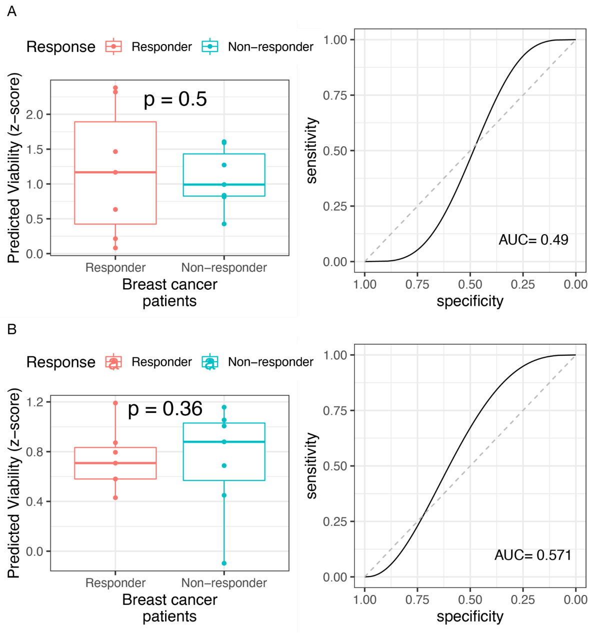


**Extended Figure 7D. PERCEPTION vs published state-of-the-art bulk response models in breast cancer clinical trial.** *Stratification power of PERCEPTION vs published state-of-the-art machine learning response models trained* ***(A)*** *only on bulk-expression (Tsherniak et al. 2017) and* ***(B)*** *PERCEPTION models that are not tuned on SC-expression. In both panels, the models*

*were generated deterministically with seed=1 during the sampling of training and test sets. The plot on the left of each panel presents the PERCEPTION predicted viability in responders vs. non-responders. The plot on right is a ROC plot depicting the prediction power (sensitivity and specificity) of the predicted viability to stratify responders vs. non-responders. The area under this curve is provided at the right corner and denotes overall model prediction power. The area under the dashed diagonal line denotes a random-model performance.*

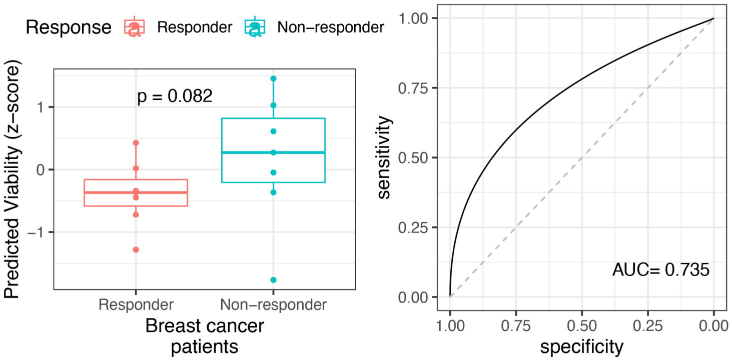


**Extended Figure 7E:** *Here, we used average SC viability instead of clone level viability to stratify responders vs. non-responders of the combination therapy arms in the FELINE clinical trial.*

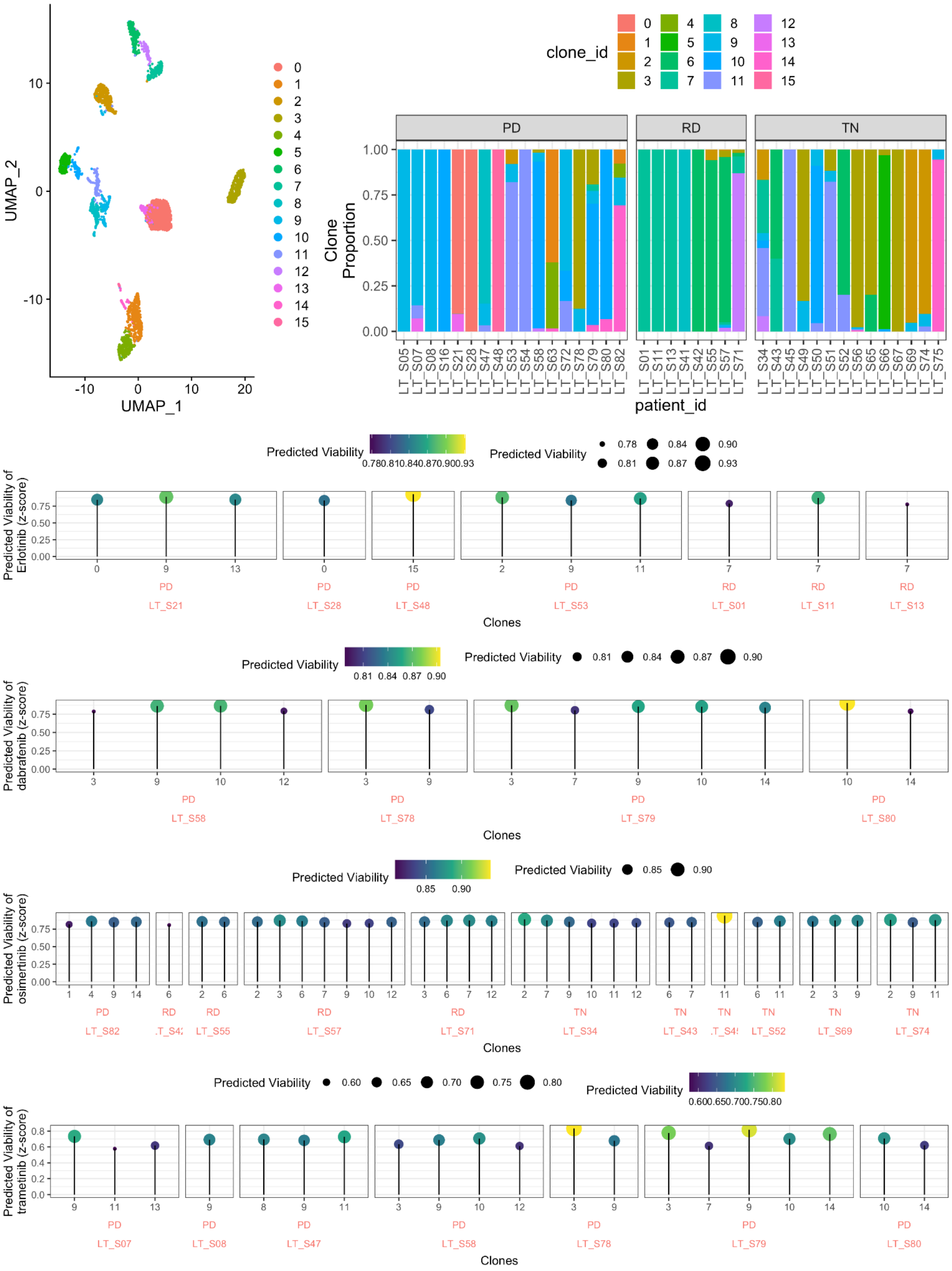


**Extended Figure 8: Pre-processing and predicting clone level response in lung cancer patients cohort.** *(A) A UMAP of 3671 malignant cells derived from 25 patients with 26485 genes are clustered using Seurat considering the first 10 axes with the most variance. Each clone (a transcriptional cluster) output is annotated using a color where the legend is provided in the*

*right. (**B**) The proportion of these clones (y-axis) are provided in each patient (x-axis) faceted by the time point at which these biopsies are collected. (**C-F**) Predicted viability of the four tyrosine kinase inhibitors: erlotinib, dabrafenib, osimertinib, and trametinib, in respective order, is provided at a clonal level for each patient where response status is provided at the bottom of each facet.*

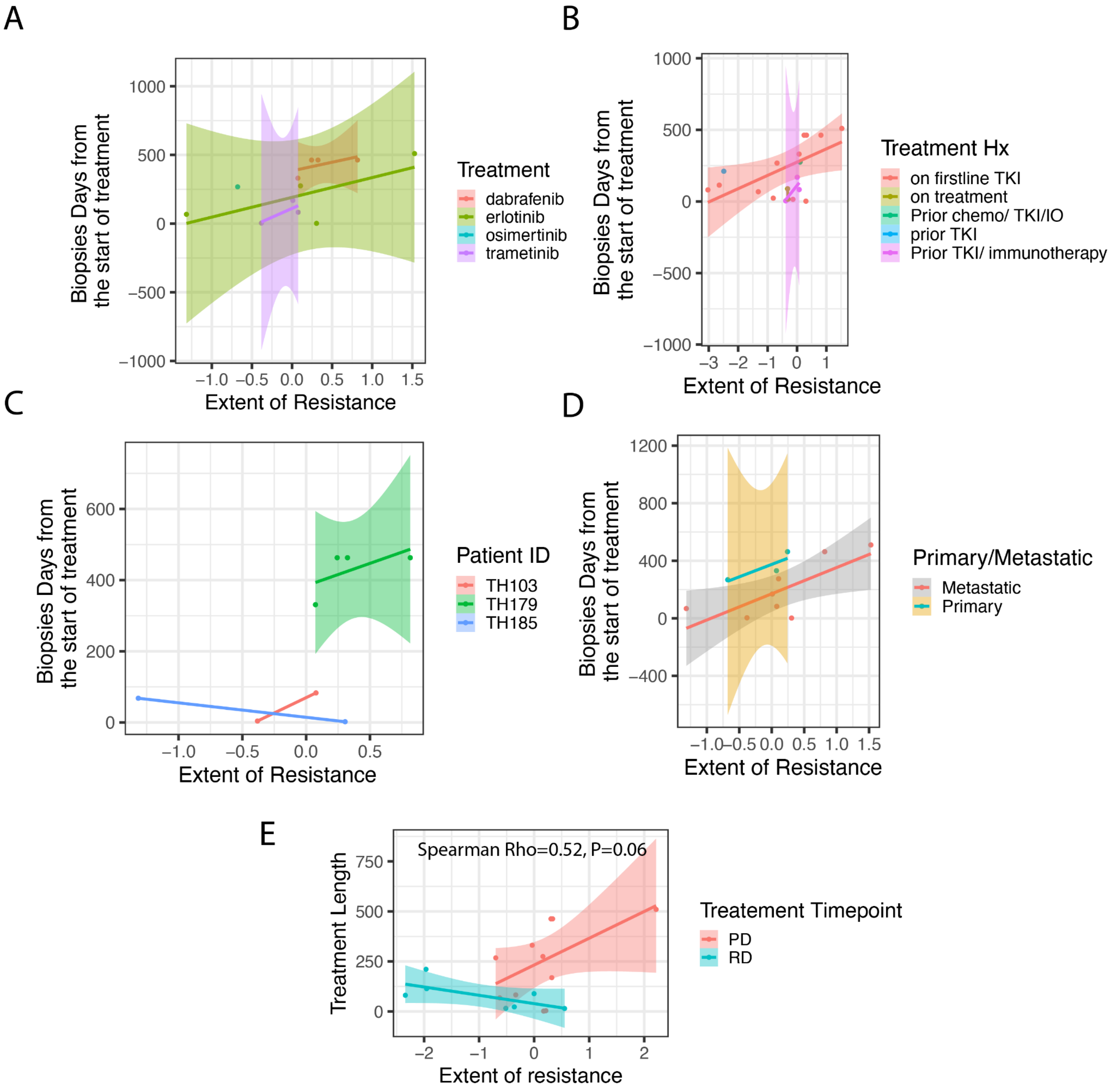


**Extended Figure 9: Correlation between the elapsed treatment time and estimated resistance holds true across different conditions.** In **A-D)**, *The extent of resistance to a*

*treatment from the baseline (x-axis) is correlated with the treatment elapsed time (Number of days from the start of the treatment before the biopsy was taken) (y-axis).* ***(A)*** *The points and line colors denote the treatment administered to the patients listed by the right legend.* ***B)*** *Color denotes prior treatment.* ***C)*** *Color denotes the patient's ID.* ***D)*** *Color denotes whether the disease is metastatic or primary at the time of biopsy.* ***E)*** *Extent of Resistance is calculated using bulk-expression of the tumor, where the increase with "Treatment Elapses time" is positive, however, insignificant and weaker than when the patient response is taken as the most-resistant clone available response.*

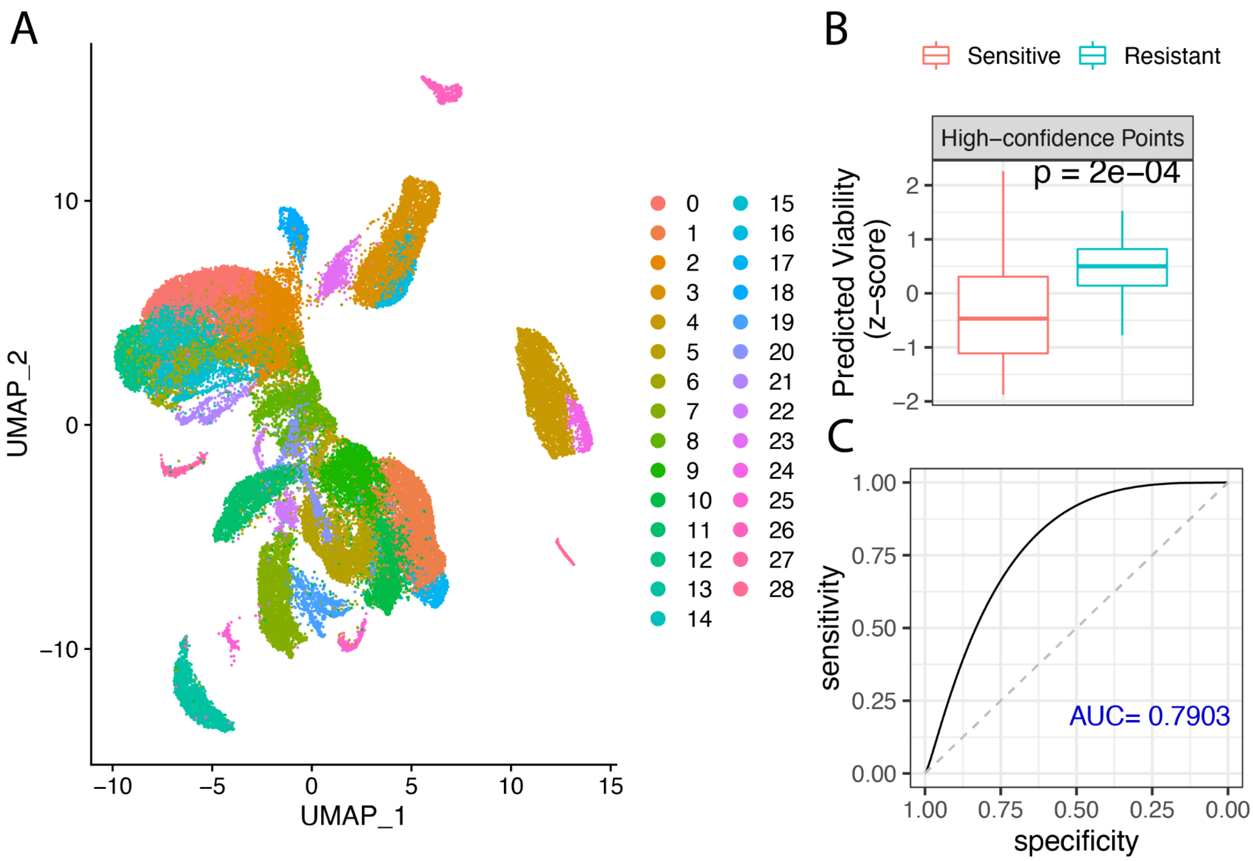


**Extended Figure 10. Prediction of monotherapy and combination response in lung cancer cell lines based on their most-resistant clone response. A)** UMap representing the clustering of 53,514 cells from 199 cell lines (~300 single cells) using their SC-expression. We noted a total of 29 clusters, where each cell had at least one single cell from four unique sub-clones (transcriptional cluster). **B)** Using this cluster information, we next provide the predicted viability, that is the viability of the most, of 21 lung cancer cell lines

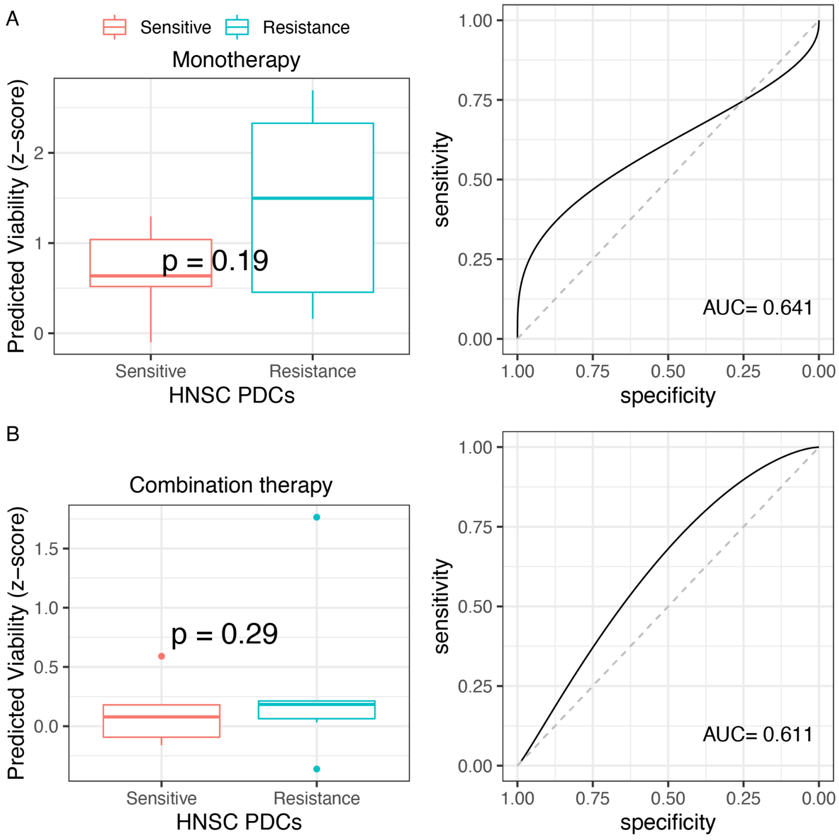


**Extended Figure 11. Prediction of monotherapy and combination response in patient-derived primary cells based on their most-resistant clone response.** *The plots present the stratification performance when* their most-resistant clone response ***(A)*** *monotherapy response and* ***(B)*** *combination response. In both the panels, the plot on the left represents the PERCEPTION predicted viability in resistance vs. sensitive cell lines. The plot on the right is a ROC plot depicting the prediction power (sensitivity and specificity) of the predicted viability to stratify resistance vs. sensitive cell lines. The area under this curve is provided at the right corner and denotes overall model prediction power. The area under the dashed diagonal line denotes a random-model performance.*

### Supplementary Notes

#### 1. SC-based PERCEPTION models prediction of monotherapy and combination response in a lung cancer cell lines screen

To study PERCEPTION in another independent screen, we tested its predictive performance in a recent drug screen in lung cancer cell lines (Nair et al. 2021). We first performed a qualitative test of the drug screen mined from (Nair et al. 2021) and only considered drugs that passed this quality control for our analysis (**Extended Figure 3A**, **Methods**). We focused on 14 FDA-approved or in clinical trials anti-cancer drugs tested in this study, for which we could build predictive PERCEPTION models. We assessed their predictive performance vs. drug screen data measured for monotherapy and two-drug combinations of these drugs across 21 lung cancer cell lines in five dosages (**Table S5, Methods**). We find that predicted viability is significantly higher in resistant vs sensitive cell lines (Top vs bottom 33% cell lines ranked by viability, Wilcoxon rank-sum P=2E-06, FC=1.53, **Extended Figure 3E**), and can stratify responders vs non-responders (ROC-AUC = 0.72). The predictive performance of the high-confidence screen (defined by subsetting the screen by only considering data points with AUC<1, **Methods**) results is considerably higher (ROC-AUC=0.88, **Figure 2C-D,** FC=1.95). The overall mean Pearson correlation between predicted vs observed viability for these drugs is 0.33; P<7.4E-09 (**Extended Figure 3B**). Observed vs predicted viability for each drug separately is provided in **Extended Figure 3C**.

We also find that the predicted combination viability is significantly higher in resistant vs sensitive cell lines (**Methods**, Wilcoxon rank-sum P=8.3E-03, FC=1.54, **Extended Figure 3E**) and can stratify the responders vs non-responders (ROC-AUC=0.69, # of resistant data points=28, # of sensitive data points=24). Like in the monotherapies case, the observed effect size is considerably higher when considering only high-confidence screen results (P<8.8E-03, FC=1.77, **Figure 2E-F**, ROC-AUC=0.87). Taken together, these results indicate the ability of PERCEPTION models to predict single and combination therapies in independent screens without any further training.

### 2. Comparison of strategies to compute clinical response from single-cell expression

We designed four strategies to compute clinical response using single cell-based clone information and tested their performance to stratify responders vs non-response in the multiple myeloma cohort. The clinical response of a patient is determined in these strategies by computing one of the following: 1. Weighted average response: an average of response across all the clones weighted by their abundance in the tumor; 2. Unweighted average response: an average of response across all the clones; 3. Most-sensitive clone response: the response of the most-sensitive clone that is the clone with the highest response; 4. Most-resistant clone response: the response of the most-resistant clone that is the clone with the least response. The AUCs for these strategies are 0.76, 0.71, 0.68 and 0.82, respectively. This testifies that the fourth strategy to use the response of the most-resistant clone response present in the tumor best reflects the clinical response.

### 3. Data processing of breast cancer clinical trial

There are a total of 34 patients with SC-expression profiles of their tumors (Griffiths et al. 2021). These patients have samples collected at different time points during their treatment, 1) collected at the time of screening (S), 2) on day 14 (M) and 3) on day 180 at the end of the trial (E). Here we show you the 36 transcription clusters identified in all 65 samples post-filtering and data processing (**Extended Figure 7A-B)**. Only the cluster annotations of patients with clinical response information are considered in our analysis (**Table S7**). We used only the treatment-naive samples (S) in Arm B and C for the stratification analysis (**Figure 5B-C, Extended Figure 7C**). We used patients with paired samples at time points S and E to study the change in response post-treatment (**Figure 5D**). The GEO downloads included only tumor cells, as a result, our analysis is limited to tumor cells only. **Extended Figure 7B**, shows the clonal distribution in each sample processed, all sub-clones which represent <5% of the cells in the sample are excluded in our analysis. **Extended Figure 7C** represents the clone level predicted response in the sample collected at day 0 in combination arms B and C. When we ignored the clonal information and predicted the patient response as the average viability across all single cells in a patient (the strategy used for cell lines and PDCs), the ability to stratify responders from non-responder diminished slightly (AUC= 0.735, **Extended Figure 7D)**.

#### 4. Data processing of in Lung cancer patients cohort and clone-level killing

The (Maynard et al. 2020) cohort is a lung cancer cohort with scRNA-seq profiles for 24 patients. We again clustered the patients with default Seurat parameters where the clusters are provided in **Extended Figure 8A**. There are a total of 16 clusters where the cluster distribution is provided in **Extended Figure 8B.** We next computed the average expression of each clone in each patient by computing the average expression across all the cells associated with the cluster in a patient and performed a rank normalization. The steps of pre-processing are precisely like the other two clinical cohorts. We next predicted treatment response using PERCEPTION of the following four drugs for each clone: erlotinib (a 1$^{st}$ generation EGFR inhibitor, **Extended Figure 8C**), dabrafenib (a BRAF inhibitor, **Extended Figure 8D**), osimertinib (3rd generation EGFR inhibitor, **Extended Figure 8E**), and trametinib (a MEK inhibitor, **Extended Figure 8F**). We next tested and found that when the analysis is repeated using bulk-expression of the tumor, the increase in *Extent of Resistance* of patients with “Treatment Elapses time” is positive, however insignificant, and lower than when the patient response is taken as the most-resistant clone available response.

#### 5. Taking cell lines response as the response observed in the most-resistant available clone

Initially, we use the average response of the single cells representing a cell line as its drug response. However, the response of the most-resistant clone in the tumor of a patient best reflected the clinical response. In this section, we present the performance when we take the cell lines response as the response observed in the most-resistant available clone to stratify resistant vs sensitive cell lines. To this end, we first clustered the 200 cell lines via Seurat using uniform parameters used across the study noting 29 clusters and with four clusters per cell line (**Extended Figure 10A**). When we considered the predicted viability of the most resistant-clone present in the cell lines as its predicted viability, we first observed that the predicted viability of sensitive cell lines is significantly lower than resistant ones (**Extended Figure 10B**) and this score could stratify the resistant vs sensitive lung cancer cell lines with an AUC of 0.79. This is stratification performance is markedly lower than the strategy where we considered a cell line response as the mean across single cells (AUC=0.89, **Extended Figure 10C**).

We repeated this process for the head and neck PDC cell lines. The transcriptional cluster/clonal information was obtained from (Suphavilai et al. 2020). For monotherapy treatments, similar to the average SC-response, the predicted viability is significantly higher in resistant vs. sensitive cell lines (top 40% vs. bottom 40% cell lines ranked by viability, N=8 each), with a ROC-AUC of 0.64 (**Extended Figure 11A**). The predicted viability over the 20 (monotherapy, cell-line) combinations (4 monotherapies x 5 cell-lines) is correlated with the observed viability (Pearson R=0.41; P<0.08). This performance diminished in combination treatments (**Extended Figure 11B**), with a ROC-AUC of 0.61 (**Extended Figure 11B**) which dropped from 0.86 (**Figure 3D**) in average SC-response. Even the predicted viability of combination treatment across the 15 (combination, therapy) cell-line pairs (3 combinations x 5 cell lines) is not highly correlated with the observed viability (Pearson R=0.34; P<0.213).

### References


Griffiths JI, Chen J, Cosgrove PA, et al. Serial single-cell genomics reveals convergent subclonal evolution of resistance as patients with early-stage breast cancer progress on endocrine plus CDK4/6 therapy. Nature Cancer 2021; 2(6): 658-671.

Maynard A, McCoach CE, Rotow JK, et al. Therapy-induced evolution of human lung cancer revealed by single-cell RNA sequencing. Cell 2020; 182(5):1232-1251.

Nair NU, Greninger P, Friedman A, Amzallag A, et al. A landscape of synergistic drug combinations in non-small-cell lung cancer. bioRxiv. 2021 [cited 2022 Jan 6]. p. 2021.06.03.447011. Available from: https://www.biorxiv.org/content/10.1101/2021.06.03.447011v1.abstract

Suphavilai C, Chia S, Sharma A, et al. Predicting heterogeneity in clone-specific therapeutic vulnerabilities using single-cell transcriptomic signatures. 2020; bioRxiv https://www.biorxiv.org/content/10.1101/2020.11.23.389676v1?rss=1

Tsherniak A, Vazquez F, Montgomery PG, et al. Defining a cancer dependency map. Cell 2017; 170(3): 564-576.
